# Zika virus infection as a cause of congenital brain abnormalities and Guillain-Barré syndrome: systematic review

**DOI:** 10.1101/073098

**Authors:** Fabienne Krauer, Maurane Riesen, Ludovic Reveiz, Olufemi T Oladapo, Ruth Martínez-Vega, Teegwendé V. Porgo, Anina Haefliger, Nathalie J Broutet, Nicola Low, for the WHO Zika Causality Working Group

## Abstract

**Background:** The World Health Organization stated in March 2016 that there was scientific consensus that the mosquito-borne Zika virus was a cause of the neurological disorder Guillain-Barré syndrome and of microcephaly and other congenital brain abnormalities, based on rapid evidence assessments. Decisions about causality require systematic assessment to guide public health actions. The objectives of this study were: to update and re-assess the evidence for causality through a rapid and systematic review about links between Zika virus infection and a) congenital brain abnormalities, including microcephaly, in the foetuses and offspring of pregnant women and b) Guillain-Barré syndrome in any population; and to describe the process and outcomes of an expert assessment of the evidence about causality.

**Methods and findings:** The study had three linked components. First, in February 2016, we developed a causality framework that defined questions about the relationship between Zika virus infection and each of the two clinical outcomes in 10 dimensions; temporality, biological plausibility, strength of association, alternative explanations, cessation, dose-response, animal experiments, analogy, specificity and consistency. Second, we did a systematic review (protocol number CRD42016036693). We searched multiple online sources up to May 30, 2016 to find studies that directly addressed either outcome and any causality dimension, used methods to expedite study selection, data extraction and quality assessment, and summarised evidence descriptively. Third, a multidisciplinary panel of experts assessed the review findings and reached consensus on causality. We found 1091 unique items up to May 30, 2016. For congenital brain abnormalities, including microcephaly, we included 72 items; for eight of 10 causality dimensions (all except dose-response relationship and specificity) we found that more than half the relevant studies supported a causal association with Zika virus infection. For Guillain-Barré syndrome, we included 36 items, of which more than half the relevant studies supported a causal association in seven of ten dimensions (all except dose-response relationship, specificity and animal experimental evidence). Articles identified non-systematically from May 30-July 29, 2016 strengthened the review findings. The expert panel concluded that: a) the most likely explanation of available evidence from outbreaks of Zika virus infection and clusters of microcephaly is that Zika virus infection during pregnancy is a cause of congenital brain abnormalities including 61 microcephaly; and b) the most likely explanation of available evidence from outbreaks of Zika virus infection and Guillain-Barré syndrome is that Zika virus infection is a trigger of Guillain-Barré syndrome. The expert panel recognised that Zika virus alone may not be sufficient to cause either congenital brain abnormalities or Guillain-Barré syndrome but agreed that the evidence was sufficient to recommend increased public health measures. Weaknesses are the limited assessment of the role of dengue virus and other possible co-factors, the small number of comparative epidemiological studies, and the difficulty in keeping the review up to date with the pace of publication of new research.

**Conclusions:** Rapid and systematic reviews with frequent updating and open dissemination are now needed, both for appraisal of the evidence about Zika virus infection and for the next public health threats that will emerge. This rapid systematic review found sufficient evidence to say that Zika virus is a cause of congenital abnormalities and is a trigger of Guillain-Barré situation.

## Introduction

An “explosive pandemic of Zika virus infection” [1] in 2015 caught the world by surprise. The PanAmerican Health Organization (PAHO) and World Health Organization (WHO) published an alert about increasing numbers of reports of “congenital anomalies, Guillain-Barré syndrome, and other neurological and autoimmune syndromes in areas where Zika virus is circulating and their possible relation to the virus” on December 1, 2015 [2]. On February 1, 2016, WHO declared that the clusters of microcephaly and other neurological disorders constituted a Public Health Emergency of International Concern [3]. Microcephaly at birth is a clinical finding indicative of reduced brain volume and can include other microscopic or macroscopic brain malformations resulting from a failure of neurogenesis [4]. Infections acquired in pregnancy, like cytomegalovirus, toxoplasmosis and rubella are established causes, and the extent and type of lesions depend on gestational stage at exposure [4]. Guillain-Barré syndrome is an immune-mediated rapidly progressing ascending flaccid paralysis, which typically occurs within a month of a bacterial or viral infection, such as *Campylobacter jejuni* and cytomegalovirus [5]. As of August 3, 2016, 65 countries in the Americas, Africa, South East Asia and Western Pacific regions have reported autochthonous transmission of the mosquito-borne flavivirus Zika since 2015 and 15 of these have reported cases of congenital brain abnormalities or Guillain-Barré syndrome or both [6]. The emergency committee of the International Health Regulations recommended increased research [3] to provide more rigorous scientific evidence of a causal relationship as a basis for the global health response to the current and future outbreaks.

Unexplained clusters of rare but serious conditions require urgent assessment of causality, balancing speed with systematic appraisal, so that public health actions can be implemented to reduce exposure to the suspected cause. Astute clinicians have often highlighted the first signals of new causes of disease in case reports [7]. But case reports are very rarely accepted as sufficient evidence of causality and need to be corroborated or refuted in a variety of different study designs [8, 9] (S1 Text, S1 Figure). Bradford Hill is widely credited for his proposed framework for thinking about causality in epidemiology in 1965, which listed nine “viewpoints” from which to study associations 100 between exposure and disease (S1 Text, S1 Table) [10]. Since then others have modified and generalised the list so that it can be applied to any putative causal relationship [11] (S1 Text, p2). Bradford Hill emphasised that his viewpoints were not rules and could not prove causation beyond doubt but, taken together, the body of evidence should be used to decide whether there is any other more likely explanation than cause and effect.

The level of certainty required before judging that Zika virus is a cause of microcephaly and Guillain-Barré syndrome is contentious [12]. Most assessments have been based on rapid but non-systematic appraisals [13–15]. Based on rapid reviews, WHO has stated that there is “scientific consensus that Zika virus is a cause of microcephaly and Guillain-Barré syndrome” since March 31, 2016 [16]. On April 13, the conclusion of a narrative review was “that sufficient evidence has accumulated to infer a causal relationship between prenatal Zika virus infection and microcephaly and other severe brain anomalies” [14]. Narrative reviews can be done quickly but typically do not describe methods for searching and selecting which studies to include, for extracting data or for assessing the methodological quality of studies. Systematic reviews typically take at least six months to complete [17], but specify research questions and methods in advance so appraisal of the evidence is more transparent and gaps in evidence can be identified [18]. Evidence about the causal relationship between Zika virus infection and Guillain-Barré syndrome has not yet been assessed. We described a causality framework for Zika virus and plans for a systematic review (S1 Text), with a preliminary overview of 21 studies about microcephaly and Guillain-Barré syndrome, published up to March 4, 2016 [19]. The objectives of this study are to re-assess the evidence for causality and update the WHO position through a rapid and systematic review about links between Zika virus infection and a) congenital brain abnormalities, including microcephaly, in the foetuses and offspring of pregnant women and b) Guillain-Barré syndrome in any population; and to describe the process and outcomes of an expert assessment of the evidence about causality.

## Methods

We describe three linked components: the causality framework for Zika virus infection, the systematic reviews and the expert panel assessment of the review findings.

### Zika causality framework

In February 2016, we developed a causality framework for Zika virus infection by defining specific questions for each of 10 causality dimensions, modified from Bradford Hill’s list (S1 Text): temporality (cause precedes effect); biological plausibility of proposed biological mechanisms; strength of association; exclusion of alternative explanations; cessation (reversal of an effect by experimental removal of, or observed decline in, the exposure); dose-response relationship; experimental evidence from animal studies; analogous cause-and-effect relationships found in other diseases; specificity of the effect; and the consistency of findings across different study types, populations and times. This review covered 35 questions about congenital brain abnormalities, including microcephaly and 26 questions about Guillain-Barré syndrome. We plan in future to examine a third group of other acute neurological disorders (S1 Text, S2 Table). We also listed seven groups of co-factors, including concurrent or previous dengue virus infection that might increase the risk of an outcome in the presence of Zika virus infection [20].

### Systematic review

Our protocol was registered on March 21, 2016 in the international database PROSPERO (number CRD42016036693) [21] and structured according to recommendations from the Preferred Reporting Items for Systematic reviews and Meta-Analysis group to structure our protocol (PRISMA-P) [22]. We report the review using the PRISMA checklist [23] and highlight features that we adopted to speed up the review process [17]. Text S1 includes methods and results that are not reported here in the main text.

To report our findings, we use the term item for an individual record, e.g. a case report, surveillance report, or original research article. Some items reported different aspects of information about the same individuals or population. To avoid double counting, we organised items that reported on the same patients into groups. We chose a primary publication (the item with the most complete information) to represent the group, to which other items were linked (S4a Table, S5a Table).

### Eligibility

We included studies of any design and in any language that directly addressed any research question in the causality framework (S1 Text). We excluded reviews, commentaries, news items and journal correspondence that did not include original data but we checked their reference lists to identify other potentially relevant studies.

### Information sources and search strategy

The search strategy was designed to find data about Zika virus and its consequences from ongoing studies and non-peer reviewed sources as well as published peer-reviewed studies to benefit from commitments to data-sharing in public health emergencies [24]. We searched: PubMed, Embase and LILACS electronic databases; PAHO Zika research portal, WHO and the European Centre for Disease Prevention and Control (ECDC) websites; journal websites; preprint servers and a real time updated portal of experimental animal studies [25] (see protocol [21] and S1 Text). For the dimension addressing analogous causes of the outcomes and for co-factors, we used items identified in the searches, their reference lists and non-systematic searches. We used Endnote X7 (Thomson Reuters, Philadelphia) for reference management.

We conducted our first search from the earliest date to April 11, 2016 and updated the search on May 30 and July 29. We selected items and extracted data systematically on included items up to May 30 and report on non-systematically identified studies up to July 29, 2016.

### Study selection and data extraction

We used pre-piloted structured forms in the online database Research Electronic Data Capture (REDCap, Vanderbilt University, Nashville). To speed up study selection we screened titles, abstract and full texts by liberal accelerated screening [17] (S1 Text) and for data extraction, one reviewer extracted data and a second reviewer checked the extracted data. Discrepancies were resolved by discussion or by a third reviewer. We did not specify a single primary outcome because the number of causality dimensions and questions was too broad [17]. The data to be extracted differed according to the study design and the question(s) addressed (S1 Text, p8 and S2 Table). We used case definitions and laboratory diagnostic test interpretations as reported by study authors. Basic research studies were too diverse to allow consistent numerical data extraction so we summarised findings descriptively.

### Synthesis of findings and assessment of methodological quality

We tabulated study level data and available data about clinical presentations from case reports, case series, cross-sectional studies, case-control studies and cohort studies. We assessed methodological quality for these designs using shortened checklists from the National Institute of Health and Clinical Excellence [26] and using reviewers’ summaries of strengths and weaknesses of other study designs. Each reviewer recorded an overall judgement of each study to indicate whether the findings did or did not provide support for the causality dimension being assessed. Two reviewers reached consensus by discussion or adjudication by a third reviewer. We assigned a judgement of sufficient evidence about a causality dimension if the reviewers’ assessments were supportive for at least half of the specific questions. We appraised the body of evidence according to the domains of the Grading of Research Assessment Development and Evaluation (GRADE) tool as suggested for urgent health questions [27], but did not apply upgrading or downgrading because these concepts could not be applied consistently across the range of study designs.

### Expert panel

The WHO Zika Research Working Group convened an expert panel of 18 members with specialist knowledge in the fields of epidemiology and public health, virology, infectious diseases, obstetrics, neonatology and neurology (members of both groups listed at the end of the article). In a series of online web and telephone conferences between April 18 and May 23, 2016, we presented our approach to the assessment of causality in epidemiological studies, the questions in our causality framework, the methods and findings of the systematic review and our synthesis of evidence for each set of clinical outcomes. We discussed these topics with the experts during the conferences and followed up through email discussions between web conferences. After the conferences and email consultation we drafted summary conclusions about the most likely explanation for the reported clusters of cases of microcephaly and Guillain-Barré syndrome. The expert panel members discussed these summaries to reach consensus statements that update the WHO position.

## Results

Figure 1 shows the timeline of the systematic review process and expert panel deliberations.

**Figure 1.**
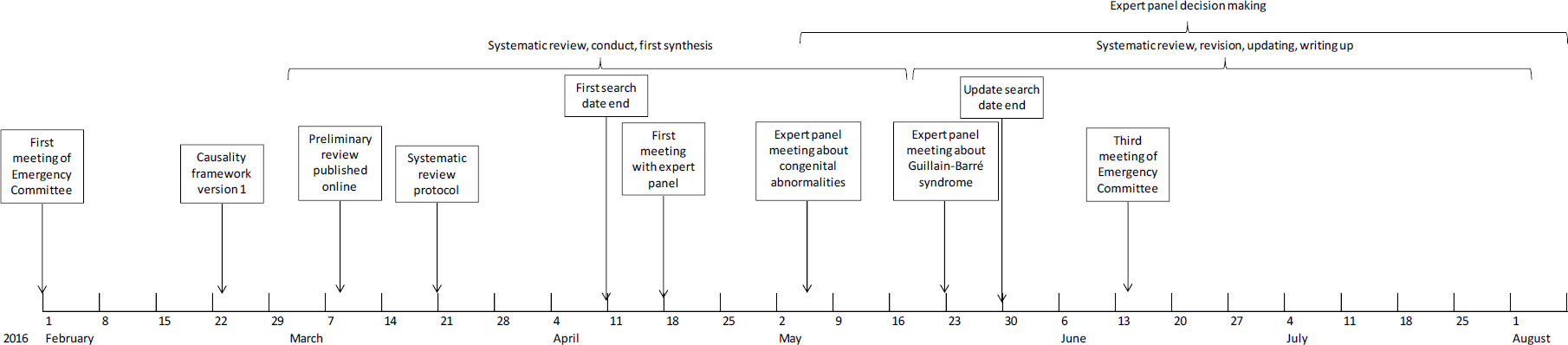
Timeline of Zika causality review, 1st February to August 2016. A Public Health Emergency of International Concern was announced on 1st February 2016 in response to clusters of microcephaly, Guillain-Barré syndrome and other neurological disorders.

We found 1091 unique items, published from 1952 to May 30, 2016 (S2 Figure, S3 Table). Most excluded items were reviews or editorials and commentaries (44%, n=479) or were articles about Zika virus that were not related to any of the causality dimensions (26%, n=282). We included 106 items from 87 groups (Table 1), of which 83% were published in 2016.

Table 1 shows the study designs and causality dimensions addressed by the included studies up to May 30, 2016. For both outcomes, the majority of items were clinical individual level case reports, 218 case series or population level surveillance data.

**Table 1.** Summary of included items according to outcome, study design and causality dimension

|  | Congenital abnormalities |  | Guillain-Barré syndrome |  |
| --- | --- | --- | --- | --- |
|  | N | % | N | % |
| **Type of study** |  |  |  |  |
| Case report | 9 | 12.5 | 9 | 25 |
| Case series | 22 | 30.6 | 5 | 13.9 |
| Case-control study | 0 | 0 | 1 | 2.8 |
| Cohort study | 1 | 1.4 | 0 | 0 |
| Cross-sectional study | 2<sup>a</sup> | 2.8 | 0 | 0 |
| Ecological study/outbreak report | 5 | 6.9 | 19 | 52.8 |
| Modelling study | 2 | 2.8 | 0 | 0 |
| Animal experiment | 18 | 25 | 0 | 0 |
| In vitro experiment | 10 | 13.9 | 0 | 0 |
| Sequence analysis and phylogenetics | 3 | 4.2 | 2 | 5.6 |
| **Total items** | **72** | **100** | **36** | **100** |
| **Causality dimension<sup>b</sup>** |  |  |  |  |
| Temporality | 21 | 36.2 | 26 | 83.9 |
| Biological plausibility | 25 | 43.1 | 4 | 12.9 |
| Strength of association | 3 | 5.2 | 2 | 6.5 |
| Alternative explanation | 18 | 31 | 6 | 19.4 |
| Cessation | 2 | 3.4 | 6 | 19.4 |
| Dose-response relationship | 0 | 0 | 0 | 0 |
| Experiment | 20 | 34.5 | 0 | 0 |
| Analogy | NA | NA | NA | NA |
| Specificity | 0 | 0 | 0 | 0 |
| Consistency | NA | NA | NA | NA |
| **Total groups<sup>c</sup>** | **58** |  | **31** |  |

<sup>a</sup> One cross-sectional study studied human participants and one studied monkeys;
<sup>b</sup> A group of items could contribute to more than one causality dimension, so totals do not sum to 100%;
<sup>c</sup> Two items contribute to both topics.
Abbreviations: NA, not applicable; evidence about analogous conditions was not searched systematically; the dimension of consistency used information in items included for all other causality dimensions.

### Congenital brain abnormalities

A total of 72 items belonging to 58 groups addressed questions related to congenital brain abnormalities up to May 30, 2016 [16, 25, 28–99]. Table 2 summarises the findings of the clinical characteristics of 278 mother-infant pairs described in case reports, case series without control groups, one cross-sectional study and one cohort study. Table 3 summarises the assessment for each causality dimension and S4a Table provides an extended description of study findings.

**Table 2.** Geographic, clinical and microbiological characteristics of mother-infant pairs

|  | No. with characteristic<sup>a</sup> | No. evaluated in the article<sup>a</sup> | %<sup>b</sup> |
| --- | --- | --- | --- |
| **Total with congenital abnormalities or adverse pregnancy outcomes** | 278 | 278 | 100 |
| **Country of infection** |  |  |  |
| Brazil | 242 | 278 | 87.1 |
| Cabo Verde | 2 | 278 | 0.7 |
| Colombia | 2 | 278 | 0.7 |
| French Polynesia | 19 | 278 | 6.8 |
| Martinique | 1 | 278 | 0.4 |
| Panama | 4 | 278 | 1.4 |
| Travellers returning from the Americas | 8 | 278 | 2.9 |
| **Pregnancy outcome** |  |  |  |
| Miscarriage | 7 | 278 | 2.5 |
| Intrauterine death or stillbirth | 3 | 278 | 1.1 |
| Termination of pregnancy | 15 | 278 | 5.4 |
| Neonatal death | 9 | 278 | 3.2 |
| Alive, still in utero | 8 | 278 | 2.9 |
| Live birth | 236 | 278 | 84.9 |
| **Time point of presumed exposure (symptoms)** |  |  |  |
| 1<sup>st</sup> trimester | 81 | 117 | 69.2 |
| 2<sup>nd</sup> trimester | 28 | 117 | 23.9 |
| 3<sup>rd</sup> trimester | 8 | 117 | 6.8 |
| **Exposure assessment in the mother** |  |  |  |
| Zika virus (ZIKV) related clinical symptoms | 180 | 265 | 67.9 |
| ZIKV positive in any test (serology/PCR/IHC) | 36 | 41 | 87.8 |
| ZIKV positive in any test before the outcome | 19 | 36 | 52.8 |
| ZIKV IgM positive (serum) | 3 | 7 | 42.9 |
| ZIKV IgG positive (serum) | 3 | 3 | 100.0 |
| ZIKV PRNT positive (serum) | 4 | 4 | 100.0 |
| ZIKV RT-PCR positive (serum) | 3 | 7 | 42.9 |
| ZIKV RT-PCR positive (urine) | 1 | 5 | 20.0 |
| ZIKV RT-PCR positive (amniotic fluid) | 9 | 12 | 75.0 |
| DENV IgG positive | 17 | 28 | 60.7 |
| **Exposure assessment in the foetus/newborn** |  |  |  |
| ZIKV positive in any test (serology/PCR/IHC) | 74 | 75 | 97.4 |
| ZIKV IgM positive (serum) | 30 | 34 | 88.2 |
| ZIKV IgG positive (serum) | 4 | 4 | 100.0 |
| ZIKV PRNT positive (serum) | 2 | 2 | 100.0 |
| ZIKV RT-PCR positive (serum) | 2 | 34 | 5.9 |
| ZIKV RT-PCR positive (brain tissue) | 6 | 6 | 100.0 |
| ZIKV RT-PCR positive (other tissue) | 6 | 11 | 54.5 |
| ZIKV RT-PCR positive (placenta/product of conception) | 7 | 8 | 87.5 |
| ZIKV RT-PCR positive (CSF) | 26 | 26 | 100.0 |
| ZIKV IHC positive (brain) | 4 | 5 | 80.0 |
| ZIKV IHC positive (other tissue) | 2 | 7 | 28.6 |
| ZIKV IHC positive (placenta/product of conception) | 3 | 4 | 75.0 |
| DENV IgG positive | 1 | 34 | 2.9 |
| **Outcome assessment** |  |  |  |
| Clinical microcephaly | 244 | 267 | 91.4 |
| Imaging confirmed brain abnormalities | 205 | 213 | 96.2 |
| Intrauterine growth restriction | 10 | 35 | 28.6 |
| Ocular disorders | 49 | 116 | 42.2 |
| Auditory disorders | 3 | 24 | 12.5 |
| Abnormal amniotic fluid | 6 | 33 | 18.2 |

<sup>a</sup> The denominator for each characteristic is the number of cases for which data were available;
<sup>b</sup> Column percentages shown for country of infection, pregnancy outcome and time point of exposure; row percentages for all other variables;
Abbreviations: CSF, cerebrospinal fluid; DENV dengue virus; IHC, immunohistochemistry; Ig, immunoglobulin; PRNT, plaque reduction neutralisation test; RT-PCR, reverse transcriptase PCR; ZIKV, Zika virus.

**Table 3.** Summary of reviewers’ assessments of evidence about Zika virus infection and congenital abnormalities, by causality dimension

| Causality dimension<sup>a</sup> | Evidence summary |
| --- | --- |
| **Temporality** | Total, 35 items in 21 groups reviewed. Reviewer assessments found sufficient evidence for all 3 questions of an appropriate temporal relationship between Zika virus (ZIKV) infection and the occurrence of congenital abnormalities, including microcephaly. The period of exposure to ZIKV was most likely to be in the first or early second trimester of pregnancy. |
| **Biological plausibility** | Total, 28 items in 25 groups reviewed. Reviewer assessments found sufficient evidence for 6 of 7 questions that address biologically plausible mechanisms by which ZIKV could cause congenital abnormalities. |
| **Strength of association** | Total, 7 items in 3 groups reviewed. Reviewer assessments found sufficient evidence of a strong association between ZIKV infection and congenital abnormalities for 2 of 2 questions. At the population level, there is strong evidence of an association. At the individual level, the effect size was extremely high, although imprecise, in 1 study and is likely to be high in the other study when follow-up is complete. |
| **Exclusion of alternative explanations** | Total, 28 items of 18 groups reviewed. Reviewer assessments found sufficient evidence at the individual level that alternative explanations have been excluded for 3 of 7 questions; no other single explanation could have accounted for clusters of congenital abnormalities. The evidence about other exposures could not be assessed because of an absence of relevant studies. |
| **Cessation** | Total, 6 items in 2 groups reviewed. Reviewer assessments found sufficient evidence for 1 of 3 questions. In two states of Brazil and in French Polynesia cases of congenital abnormalities decreased after ZIKV transmission ceased. Evidence for the other questions could not be assessed because no relevant studies were identified. |
| **Dose-response relationship** | This dimension could not be assessed because of an absence of relevant studies. |
| **Animal experiments** | Total, 20 items reviewed. Reviewers assessments found evidence from animal experimental studies for all 4 questions that supports a causal link between ZIKV and congenital abnormalities. Inoculation with ZIKV of pregnant rhesus macaques and mice can result in foetal abnormalities, viraemia and brain abnormalities. Experiments to induce viral replication after inoculation of ZIKV intracerebrally and at other sites in a variety of animal models have produced mixed results. |
| **Analogy** | Selected studies reviewed. There are analogies with the well-described group of TORCH infections. Microcephaly has been described following the flavivirus West Nile virus (WNV) infection in pregnancy but not DENV. Evidence was not reviewed systematically. |
| **Specificity** | No items reviewed. We did not find any studies that identified congenital abnormalities that were found following Zika virus infection in pregnancy but not in other congenital infections. The studies included described a wide range of abnormalities on clinical and neuroimaging examinations. Many of the abnormalities described are also found in other congenital infections, but with a different pattern. |
| **Consistency** | For 3 of 4 questions, the evidence assessed was consistent. By geographical region, maternal exposure to ZIKV has been associated with the occurrence of congenital abnormalities in all three regions where ZIKV has circulated since 2007. By study design, the association between ZIKV infection and congenital abnormalities has been found in studies at both individual and population level and with both retrospective and prospective designs. By population group, ZIKV infection has been linked to congenital abnormalities in both women resident in affected countries and in women from non-affected countries whose only possible exposure to ZIKV was having travelled in early pregnancy to an affected country. The evidence according to ZIKV lineage is inconsistent because an association between ZIKV and congenital abnormalities has only been reported from countries with ZIKV of the Asian lineage since 2013. |

<sup>a</sup> Questions for each causality dimension are in S1 Text, S2 Table.
Abbreviations: DENV, dengue virus; TORCH, Toxoplasmosis, Rubella, Cytomegalovirus, Herpes simplex virus; WNV, West Nile virus; ZIKV, Zika virus.

### Temporality

Thirty-five items [37–44, 46–50, 52–54, 56, 58, 59, 64, 67, 72, 74, 75, 79–82, 84, 86, 91, 92, 94–96] in 21 groups addressed three questions about this dimension (Table S4a). Overall, 67.9% (180/265) of women with clinical data available reported Zika virus symptoms during pregnancy (Table 2). The temporal sequence of confirmed Zika virus infection preceding a diagnosis of microcephaly was only available in a small proportion of pregnant women because many case reports were published before laboratory confirmation testing was available. Of the 36 mothers with laboratory confirmed Zika virus infection (serology and/or reverse transcriptase PCR, RT-PCR) 19 (52.8%) were confirmed before the detection of foetal malformations or the occurrence of miscarriage [47, 49, 52, 59, 74]. The most recent studies show detailed timelines of laboratory confirmation of recent infection followed by in utero neuroimaging evidence of brain abnormalities and subsequent birth with microcephaly [49] or termination of pregnancy and confirmation of foetal infection [59]. The most likely time point of exposure was the first trimester or the early second trimester, based on individual case reports and three statistical modelling studies [54, 56, 67]. At the population level, epidemic curves of possible cases with Zika virus illness increased in parallel to reported cases of microcephaly with a time gap of 30 to 34 weeks in two states of Brazil (Pernambuco and Bahia) [64, 67] (S1 Text, p9, S3 Figure).

### Biological plausibility

Twenty-eight items [36, 37, 41, 43–45, 47, 48, 51, 58, 59, 61–65, 68, 70, 71, 74, 76, 78, 81, 83, 85, 87, 91, 97–99] in 25 groups addressed seven questions about this dimension of causality (S4a Table). The studies suggest several biologically plausible effects of Zika virus transmission in utero. Detailed investigations from one report about a woman found Zika virus by RT-PCR in the serum of the woman with normal foetal ultrasound at 13 weeks [59]. Four weeks later, ultrasound showed a decrease in head circumference and other brain abnormalities and the pregnancy was terminated. The isolated viral particles from the brain were capable of replication in cell culture, but particles isolated from other tissues were not. Zika virus RNA was also found in foetal brain tissue in three other studies [41, 43, 45]. Basic research experiments have also found evidence that Zika virus from both the African and the Brazilian (Asian) lineages replicates in different types of neural progenitor cells [51, 76, 97]. The phosphatidylserine-sensing receptor tyrosine kinase AXL is a potential entry point into human cells; AXL has also been found to be expressed in developing human cerebral cortex tissue [36, 71]. *In vitro* studies using neural progenitor cells (NPCs) and cerebral organoids show that Zika virus replicates in neural tissue and can disturb the cell cycle and lead to apoptosis [51, 76, 83, 87, 97]. These findings suggest a teratogenic effect of Zika virus on the developing brain in which dysregulation of cell division and apoptosis during embryonal and/or foetal development contribute to the pathogenic effects.

### Strength of association

We reviewed seven items [49, 53, 54, 56, 67, 84, 92] in three groups up to May 30, 2016 for this dimension (S4a Table). Two published studies suggest that the association between Zika virus infection in pregnancy and congenital brain abnormalities is likely to be very strong [49, 54]. In Rio de Janeiro, investigators modified an ongoing study of women with rash in pregnancy [49]. They compared 72 women with positive RT-PCR results for Zika virus with 16 women with other causes of rash. Follow-up and assessment of the outcome seems to have been more intensive in women with Zika virus infection than those without. Of 42 Zika-infected women with one or more ultrasound scans, 12 (29%) had abnormal scans. All 16 women without Zika virus infection were reported to have had one normal routine scan, but no follow up data were reported. The authors did not calculate a risk ratio but the descriptive preliminary data suggest that Zika virus infection was associated with a marked increase in the risk of a wide range congenital abnormalities. In French Polynesia, investigators re-constructed a hypothetical cohort of pregnant women from different sources of data, including eight retrospectively identified cases of microcephaly. They estimated that the risk of microcephaly would be 53.4 times (95% confidence interval 6.5–1061.2) higher in women with Zika virus infection than in uninfected women if all infections had occurred in the first trimester. The statistical modelling and assumptions were clearly described, but the estimate was obtained from indirect data sources and the confidence intervals are very wide. A case-control study in Brazil, completed after May 30 was also identified[100].

At population level, analyses of data at the level of the state in Brazil showed a positive correlation between case reports of Zika-like illness per 100,000 population and cases of microcephaly per 100,000 live births [56]. A separate analysis of these data showed a higher prevalence of microcephaly in 15 states that had reported Zika virus cases (2.8 per 10,000 live births) than in four states with no reported cases (0.6 per 10,000 live births) [53], corresponding to a prevalence ratio of 4.7 (95% CI 1.9-13.3). The authors acknowledge potential under-reporting before surveillance was enhanced in 2015-2016, but the prevalence of microcephaly in the two worst affected states was still more than twice the level of a previous estimate from 1995-2008 of 5.1 per 10,000 births.

### Exclusion of alternative explanations

Twenty-eight items [37–46, 48–50, 52, 58, 59, 72, 75, 79–82, 85, 86, 91, 94–96] in 18 groups addressed three of six pre-specified categories of alternative explanations (S4a Table). From these assessments, no alternative single infectious cause could have resulted in large clusters of cases of microcephaly in different places. Sporadic cases with syphilis or HSV were found, but most mothers or infants were negative or seroconverted (negative IgM and positive IgG) for cytomegalovirus, rubella and toxoplasmosis. Acute dengue virus infection was also excluded in most studies. A small number of studies excluded maternal exposure to alcohol or medication, or genetic causes of congenital abnormalities [41, 42, 44, 58]. No study excluded exposure to environmental toxins or heavy metals.

### Cessation

We reviewed six items [53, 56, 64, 67, 84, 92] in two groups that addressed one of three questions about this dimension (S4a Table). Surveillance reports of cases of suspected Zika virus-like illness in northeastern Brazil in 2015 declined [64, 67] either due to seasonality of the vector or population immunity. Reports of microcephaly cases declined with a similar temporal pattern in Bahia state [67]. In Pernambuco state, a similar decrease in Zika cases and microcephaly notifications was observed but a dengue epidemic occurred simultaneously with case numbers exceeding reports of Zika virus illness throughout 2015 so the decline in microcephaly cases might not be attributable to the Zika outbreak alone [64] (S1 Text, S3 Figure). There is no vaccine or treatment so it cannot be shown that a deliberate intervention would reverse the trend. We did not find any data on trends in microcephaly cases in countries other than Brazil.

### Dose-response relationship

We did not find any studies that addressed this dimension of causality.

### Experiments in animals

We reviewed 20 items [25, 28–35, 55, 57, 60, 66, 69, 76, 77, 88–90, 93] that addressed four questions 328 about animal experiments (S4a Table). Studies in the 1950s-1970s shows that experimental inoculation of Zika virus resulted in illness, cerebral lesions and viral replication in the brain in some but not all species tested [28–32, 34, 35]. Some of these effects might have been enhanced by the numerous serial passaging and subsequent viral adaptation of the original Ugandan Zika strain MR766 and the choice of genetically susceptible animal models. Wild monkeys with Zika virus, captured in Ethiopia, were also found to have degenerative brain lesions, but these lesions were not necessarily caused by the virus [33]. From 2000 onwards, animal studies have shown evidence of neurotropism in immunocompromised young and adult mice (A129, AG129, SCID, Ifnar) that lack are vulnerable to virus infections and in foetal or infant (suckling) immunocompetent mice (C57, BALB/c) [55, 77, 88], but not in adult immunocompetent mice (129 Sv/Ev, CD1, C57) [57, 60]. Real time reports are documenting studies of Macaque monkeys, experimentally infected with a Brazilian strain and a French Polynesian strain of Zika virus (both are Asian lineage) during pregnancy [25]. High and persisting viraemia was observed in one animal. The infant did not have clinical microcephaly at delivery and brain tissue was negative for viral RNA, but some foetal tissues were positive. Inoculation of pregnant immunocompromised mice showed that Zika virus could cross the placenta and killed most embryos. The remaining foetuses showed significant growth restriction but not microcephaly [90].

### Analogy

The link between clusters of babies born with microcephaly and an earlier outbreak of Zika virus infection in Brazil is analogous to an astute clinician’s description in 1941 of a cluster of babies with congenital cataracts, microphthalmia and other abnormalities linked to an outbreak of rubella seven months before in Australia [101]. Some of the clinical features described in infants born to mothers who had Zika virus infection in pregnancy are similar to the consequences of other congenital infections, including rubella. Cytomegalovirus and toxoplasmosis can both cause microcephaly, intracranial calcification and ocular and auditory defects [102] (cited in [50]). Two cases of microcephaly were reported amongst 72 women infected with the neurotropic flavivirus West Nile virus infection in pregnancy [103]. A review of 30 studies of dengue virus infection in pregnancy found evidence of vertical transmission but did not mention microcephaly or other congenital brain abnormalities as possible complications [104].

### Specificity of association

We did not find any studies that described neuroimaging or clinical features found only in association with Zika virus infection. A preliminary article described qualitative similarities and differences compared with other congenital infections [50] and several uncontrolled case series described the spectrum of neurological and other physical abnormalities *in utero* and at birth [39, 49, 58, 105].

### Consistency

We assessed evidence of consistency by study design, geography, sub-population and virus lineage in all included items. Findings that support Zika virus infection as a cause of congenital brain abnormalities have come from different kinds of epidemiological studies and laboratory studies in both humans and animals (S4a Table). Case reports of pregnancies affected by Zika virus have come from different parts of the Americas, the Pacific region (Table 2) and West Africa [16, 73]. The prevalence of microcephaly has not been higher than expected in all countries with Zika virus transmission, however. Congenital brain abnormalities or Zika virus infection in products of conception, diagnosed in pregnant women returning from travel to a Zika-affected country [41, 47, 59], show consistency across populations. There have been no reports of congenital brain abnormalities from countries affected by the African lineage [106]. One in vitro study found that Brazilian (Asian lineage) and African Zika strains both replicated in murine and human cell cultures and organoids [76, 83]. Rhesus macaques infected with the French Polynesian strain showed higher viraemia than macaques infected with the African lineage [66].

### Summary of quality of evidence

The body of evidence includes a wide range of study designs and populations in both humans and animals (S4b Table). Much of the evidence in humans comes from uncontrolled or ecological study designs that have inherent biases for ascertaining causal associations. Amongst the few studies that examined the strength of association, effect sizes were either very large or (in an ongoing study) expected to be very large but also imprecise. One of three comparative studies was at low risk of bias. Evidence from animal studies is, by its nature, indirect. We could not formally assess publication bias; our search strategy was wide but we found very few studies with findings that were not consistent with causality. Evidence about analogous situations was not reviewed systematically.

### Guillain-Barré syndrome

We found 35 items belonging to 31 groups that addressed questions related to Guillain-Barré syndrome [61–64, 74, 84, 107–129]. We summarise the findings according to clinical characteristics of 117 individuals diagnosed with Guillain-Barré syndrome in case reports, case series without control groups and case-control studies in Table 4. Table 5 summarises the reviewers’ assessments by causality dimension and S5a Table provides an extended description of study findings.

**Table 4.** Geographic, clinical and microbiological characteristics of people with Guillain-Barré syndrome

|  | No. with characteristic | No. evaluated | % |
| --- | --- | --- | --- |
| **Total N of cases with Guillain-Barré syndrome** | 117 | 117 | 100 |
| **Country of infection** |  |  |  |
| Brazil<sup>a</sup> | 43 | 117 | 36.8 |
| El Salvador<sup>a</sup> | 22 | 117 | 18.8 |
| French Polynesia | 42 | 117 | 35.9 |
| Haiti | 1 | 117 | 0.9 |
| Martinique | 2 | 117 | 1.7 |
| Panama | 2 | 117 | 1.7 |
| Puerto Rico | 1 | 117 | 0.9 |
| Travellers returning from the Americas | 3 | 117 | 2.6 |
| Venezuela | 1 | 117 | 0.9 |
| **Exposure assessment** |  |  |  |
| Zika virus (ZIKV) symptomatic cases | 83 | 112 | 74.1 |
| ZIKV positive in any test (serology/RT-PCR) | 53 | 53 | 100.0 |
| ZIKV IgM positive (serum) | 41 | 44 | 93.2 |
| ZIKV IgG positive (serum) | 29 | 42 | 69.0 |
| ZIKV PRNT positive (serum) | 43 | 43 | 100.0 |
| ZIKV RT-PCR positive (serum) | 3 | 49 | 6.1 |
| ZIKV RT-PCR positive (urine) | 5 | 6 | 83.3 |
| ZIKV RT-PCR positive (saliva) | 0 | 0 |  |
| ZIKV RT-PCR positive (CSF) | 1 | 3 | 33.3 |
| ZIKV culture positive (serum) | 0 | 0 |  |
| ZIKV culture positive (CSF) | 0 | 0 |  |
| DENV IgG positive | 43 | 45 | 95.6 |
| **Interval between ZIKV and Guillain-Barré syndrome symptoms, days** | Median 10, range 3-12 [113, 115, 117, 118, 124, 127]<br>French Polynesia: Median 6 (IQR 4-10) [119]<br>El Salvador: 7-15 [115] |  |  |

<sup>a</sup> Only one patient with Guillain-Barré syndrome in Brazil and none in El Salvador had laboratory confirmation of Zika virus infection;
Abbreviations: CSF, cerebrospinal fluid; DENV dengue virus; IQR, interquartile range; Ig, immunoglobulin; PRNT, plaque reduction neutralisation test; RT-PCR, reverse transcriptase PCR; ZIKV, Zika virus.

**Table 5.** Summary of reviewers’ assessments of evidence about Zika virus infection and Guillain-Barré syndrome, by causality dimension

| Causality dimension<sup>a</sup> | Evidence summary |
| --- | --- |
| 1. **Temporality** | Total, 31 studies in 26 groups. Reviewer assessments found sufficient evidence for all 3 questions of an appropriate temporal relationship between ZIKV infection and GBS. The time interval between ZIKV symptoms and onset of neurological symptoms was compatible with that of other accepted triggers of GBS. |
| 2. **Biological plausibility** | Total, 6 items in 4 groups reviewed. Reviewer assessments found sufficient evidence for 2 of 3 questions about biologically plausible mechanisms by which ZIKV could trigger the immune-mediated pathology of GBS. There is evidence that supports a role for molecular mimicry, a proposed mechanism of autoimmunity, which has been reported in *Campylobacter jejuni*-associated GBS. Direct neurotropic effects of ZIKV might also occur. |
| 3. **Strength of association** | Total 7 items in 2 groups reviewed. The reviewers assessed evidence from the ZIKV outbreak in French Polynesia as showing a strong association between ZIKV and GBS at both the individual and population level. Surveillance reports from Brazil also support an association at the population level. Preliminary results from a case-control study in Brazil suggest a similar, strong effect. |
| 4. **Exclusion of alternative explanations** | Total, 10 items in 7 groups studies reviewed. Reviewer assessments found sufficient evidence at the individual level that other infectious triggers of GBS have been excluded; no other single infection could have accounted for clusters of GBS. The evidence about other exposures could not be assessed because of an absence of relevant studies. |
| 5. **Cessation** | Total 8 items in 6 groups reviewed. Reviewer assessments found sufficient evidence for 1 of 3 questions. In one state in Brazil, four other countries in the Americas and in French Polynesia, reports of GBS decreased after ZIKV transmission ceased. Evidence for the other questions could not be assessed because no relevant studies were identified. |
| 6. **Dose-response relationship** | No relevant studies identified. |
| 7. **Animal experiments** | No relevant studies of animal models of immune-mediated neuropathology identified. Evidence about neurotropism of ZIKV summarised in S4a Table. |
| 8. **Analogy** | Evidence was not reviewed systematically; Selected studies reviewed for 2 of 3 questions. Analogous mosquito-borne neurotropic flavivirus infections have been reported in association with GBS (WNV; DENV; JEV). WNV and JEV have also been reported to be associated with direct neurotropic effects and poliomyelitis-like acute flaccid paralysis. The time lag between ZIKV symptoms and GBS symptoms is analogous to intervals reported for other infectious triggers of GBS. There is some evidence that, as for *C. jejuni*-associated GBS, molecular mimicry could be involved. |
| 9. **Specificity** | No relevant studies identified. |
| 10. **Consistency** | For 3 of 4 questions, there was sufficient evidence of consistency. By geographical region, ZIKV transmission has been associated with the occurrence of GBS in 2 of 3 regions where ZIKV has circulated since 2007. By study design, the association between ZIKV infection and GBS has been found in studies at both individual and population level. By population group, ZIKV infection has been linked to GBS in both residents of an affected country and travellers from non-affected countries whose only possible exposure to ZIKV was having travelled to an affected country. The evidence according to ZIKV lineage is unclear because an association between ZIKV and GBS has only been reported from countries with ZIKV of the Asian lineage since 2013. |

<sup>a</sup> Questions for each causality dimension are in S1 Text, S2 Table.
Abbreviations: DENV, dengue virus; GBS, Guillain-Barré syndrome; JEV, Japanese encephalitis virus; WNV, West Nile virus; ZIKV, Zika virus.

### Temporality

We included 31 items [62–64, 74, 84, 107–119, 122–124, 127–129] in 26 groups that addressed three questions about this dimension (S5a Table). A temporal association at the individual level has been shown, with symptoms of Zika virus infection reported before the onset of Guillain-Barré syndrome symptoms in cases in French Polynesia, Brazil, El Salvador, Panama, Puerto Rico and Venezuela, and in returning travellers from Haiti, Suriname and Central America. All patients with Guillain-Barré syndrome had laboratory confirmed Zika virus infection except for 42 of 43 in Brazil and all those in El Salvador. The intervals between Zika virus and neurological symptoms delays of three to 12 days [115, 118, 119] are consistent with a post-infectious autoimmune mechanism [5]. In one ecological study in Bahia, Brazil, the lag between the epidemic peaks of cases with acute exanthematous illness and Guillain-Barré syndrome was five to nine weeks; the authors concluded that the actual delay might be shorter because the surveillance data recorded the date of hospitalisation rather than the onset of symptoms [84].

At the population level, 11 countries in Latin America (Brazil, Colombia, El Salvador, French Guiana, Honduras, Venezuela, Suriname) and the Caribbean (Dominican Republic, Jamaica, Martinique) and French Polynesia have reported an increase in Guillain-Barré syndrome cases during outbreaks of Zika virus infection. Surveillance reports show sporadic Guillain-Barré syndrome cases in association with Zika in four countries but without an increase above background level (Guadeloupe, Haiti, Panama, Puerto Rico). One study reported on surveillance data about acute flaccid paralysis in children, which is conducted routinely as part of the surveillance system for polio, in 20 island states in the South Pacific. The numbers of expected cases of acute flaccid paralysis was <1 per year in most countries because populations are small and an increase during periods of Zika virus transmission was only observed in the Solomon Islands [122].

### Biological plausibility

We reviewed six items [61, 109, 118, 119, 121, 123] in four groups that addressed two of three questions about biologically plausible mechanisms by which Zika virus could act as a trigger of Guillain-Barré syndrome (S5a Table). Anti-ganglioside antibodies, whose presence supports the clinical diagnosis of Guillain-Barré syndrome, were found in the serum of a third of patients in a case-control study in French Polynesia [119] and in one patient from Venezuela [118]. The case-control study and two *in silico* studies also provide some evidence for molecular mimicry of Zika virus epitopes and host antigens [119]. The *in silico* comparison of predicted epitopes and human antigens suggested peptide sharing between Zika virus and human proteins related to myelin/neuropathy [121] and von Willebrand Factor [61]. A direct effect of Zika virus on anterior horn cells or neurons might also be plausible. Several experimental studies with human neural stem cells and various mouse models have shown some evidence for neurotropism of Zika virus (see S4a Table).

### Strength of association

We reviewed seven items [108–111, 119, 123, 129] in two groups identified up to May 30, 2016. We found one published case-control study, which enrolled 42 cases of Guillain-Barré syndrome during the Zika outbreak in French Polynesia and compared them with two control groups, 98 patients hospitalised at the same time with non-febrile illness and 70 patients with acute Zika virus illness but no neurological symptoms [119] (S5a Table). Several alternative causes of Guillain-Barré syndrome were excluded. Evidence of Zika virus infection was much more common in Guillain-Barré syndrome cases than controls (odds ratios 59.7, 95% CI 10.4–+∞ defined as IgM or IgG positivity and 34.1, 95% CI 5.8–+∞ defined as presence of neutralising antibodies). Cases and controls were matched but there was no additional adjustment for confounding. Using the same cases and population denominators, the incidence of Guillain-Barré syndrome in French Polynesia was estimated to be 21 time higher during the Zika epidemic than in the pre-Zika period of 2009 to 2012, an attributable risk of 0.39 per 1000 py [129]. In Brazil, surveillance data showed a 19% increase in reports of Guillain-Barré syndrome cases in 2015 compared with 2014 in the country as a whole [108]. Information received after May 30 found a second case-control investigation conducted in Brazil that enrolled controls from the community and is ongoing; preliminary results suggest a similar, strong effect.

### Alternative explanations

We included ten items [109, 117, 119, 120, 123–128] in seven groups that addressed one of four categories of alternative explanations (S5a Table). In several studies, other infections that can trigger Guillain-Barré syndrome were excluded, such as *C. jejuni*, *Mycoplasma pneumoniae*, HIV, Epstein-Barr virus and herpes simplex virus. In the included studies, no single infectious trigger that would have resulted in Guillain-Barré syndrome outbreaks in multiple geographical locations was identified.

### Cessation

Eight items [63, 64, 84, 110, 111, 116, 129] in six groups addressed one of three questions about the effects of the removal of the suspected exposure (S5a Table). In surveillance reports from six countries (Brazil, Colombia, El Salvador, French Polynesia, Honduras and Suriname) the incidence of Guillain-Barré syndrome declined as reports of Zika virus infection fell. There is no vaccine or treatment so it cannot be shown that a deliberate intervention would reverse the trend.

### Dose-response relationship, experiments in animals and specificity

We did not find any studies that addressed these dimensions of causality.

### Analogy

Guillain-Barré syndrome is a para or post-infectious neurological condition that can be triggered by a range of viral and bacterial infections [5]. Clusters of cases of Guillain-Barré syndrome have been reported in association with outbreaks of *C. jejuni* gastroenteritis [130]. The incidence of Guillain-Barré syndrome estimated from studies of the outbreak in French Polynesia of 0.24 per 1000 Zika virus infections [119], is at the lower end of estimates from studies of *C. jejuni* (0.3 per 1000 [131] and 1.17 per 1000 [132]). The reported latency between gastrointestinal symptoms and onset of paralysis of approximately 9 days (range 1-23 days) [131, 133, 134] is similar to Zika virus-associated cases. Other, mosquito-borne neurotropic flaviviruses have been reported as possible triggers of Guillain-Barré syndrome in case reports and case series; dengue virus [135], West Nile virus [136], Japanese B encephalitis virus [137, 138] or yellow fever 17D vaccination [139]. An acute poliomyelitis-like flaccid paralysis, resulting from direct neural infection presumably of anterior horn cells, has also been reported as a clinical consequence of these viruses [136, 140, 141]. Putative biological mechanisms include upregulation of MHC class I and II molecules of peripheral nerve cells and subsequent immune-mediated cell destruction [142], auto-antibodies directed against heat shock proteins [143], galactocerebrosides [144] or myelin basic protein (MBP), and proliferation of MBP specific T-cells [145].

### Consistency

We assessed evidence of consistency by study design, geography, sub-population and virus lineage in all included items (S5a Table). The link between Zika virus and Guillain-Barré syndrome has been made in studies of different designs at individual and population level. Clusters of Guillain-Barré syndrome have been seen in multiple countries during epidemics of Zika virus but have not been reported in all those in which Zika virus outbreaks have occurred. Outbreaks of Guillain-Barré syndrome in which gene sequencing has been done were associated with Zika virus of the Asian lineage.

### Summary of quality of evidence

The body of evidence includes a wide range of study designs and populations in humans (S5b Table). A majority of the evidence reviewed was from uncontrolled or ecological study designs that have inherent biases for ascertaining causal associations. The only study that examined the strength of association found a very large but imprecise estimate of the effect size. This study did not have serious risks of bias. There was no evidence of indirectness. We could not formally assess publication bias but we had a broad search strategy and we did find evidence that outbreaks of Guillain-Barré syndrome have not been seen in all countries with Zika virus transmission.

### Co-factors that might act in the presence of Zika virus

We prespecified seven categories of co-factors (S1 Text, S2 Table). The most widely discussed in the studies that we reviewed was past dengue virus infection [119]. It is hypothesised that a mechanism known as antibody-dependent enhancement might be involved, when IgG antibodies against viral envelope proteins resulting from a prior infection bind to virus particles of a subsequent infection leading to enhanced replication and potentially more severe illness [146]. Evidence from *in vitro* experiments suggests cross-reactivity between dengue and Zika virus antibody responses and antibody dependent enhancement of Zika virus by dengue antibodies [146, 147]. In several of the studies that we reviewed, evidence of past dengue virus infection was reported (S1 Text, p10-11). We did not systematically review evidence for other co-factors but report additional narrative findings in S1 Text.

### WHO expert panel conclusions about causality

- The most likely explanation of available evidence from outbreaks of Zika virus infection and clusters of microcephaly is that Zika virus infection during pregnancy is a cause of congenital brain abnormalities including microcephaly;
- The most likely explanation of available evidence from outbreaks of Zika virus infection and Guillain-Barré syndrome is that Zika virus infection is a trigger of Guillain-Barré syndrome.

The expert panel recognises that Zika virus alone may not be sufficient to cause either congenital brain abnormalities or Guillain-Barré syndrome. The expert panel recognises that Zika virus alone may not be sufficient to cause either congenital brain abnormalities or Guillain-Barré syndrome. We do not know whether these effects depend on as yet uncharacterised co-factors being present; nor do we know whether dengue virus plays a part; as this is carried by the same species of mosquito and has circulated in many countries during the same period.

The panel agreed that there is sufficient evidence to recommend increasing:

- public health actions to reduce the risk of the effects of Zika virus infection in pregnancy, and to provide appropriate care and support for women who have been exposed [148];
- public health actions to reduce exposure to Zika virus for all people;
- public health actions to provide appropriate clinical care and rehabilitation and continuing care for all those with long term neurological conditions, such as acute clinical services and rehabilitation;
- surveillance and research into diagnostics, vaccines, treatments and vector control.

## Discussion

We conducted a rapid systematic review of 109 items from 87 groups up to May 30, 2016 about causal links between Zika virus infection and congenital brain abnormalities or Guillain-Barré syndrome. We found at least one study that supported a causal association between Zika virus infection and congenital brain abnormalities, including microcephaly, addressing one or more specific questions for eight of 10 causality dimensions (all except dose-response relationship and specificity) and Guillain-Barré syndrome in seven of ten dimensions (all except dose-response relationship, specificity and animal experimental evidence). There are methodological weaknesses, inconsistencies and gaps in the body of evidence for both sets of conditions. Studies found after the cut-off for our first searches did not change our conclusions, but strengthened the evidence about biological plausibility, strength of association and exclusion of alternative explanations.

### Interpretation of the review findings

The expert panel’s conclusions support causal links between Zika virus infection and congenital brain abnormalities and Guillain-Barré syndrome and address Bradford Hill’s pragmatic question, “is there any other way of explaining the set of facts before us, is there any other answer equally, or more, likely than cause and effect?” [10]. The conclusions are based on the body of evidence, which includes both the epidemiological context of unexpected clusters of different types of neurological conditions in countries that have experienced their first outbreaks of Zika virus infection and the strengths and weaknesses of a systematic review structured around 10 dimensions of causality (S4a Table, S4b Table, S5a Table and S5b Table). Empirical observations cannot “prove” causality, however [10, 149], and discussions about Zika virus and the terminology for describing its effects have been intense [12]. We use the term “a cause” rather than “the cause” because most causes of disease are just one component of a set of factors that all have to be present in the right constellation to result in the effect [150]. A cause can be identified without understanding all the other components or the complete causal mechanisms involved [149, 150]. In the case of Guillain-Barré syndrome, the infections that precede it are often referred to as “triggers” of the immune-mediated causal pathways involved in pathogenesis.

The body of evidence about Zika virus and congenital abnormalities (72 items included in the systematic review) has grown more quickly than that for Guillain-Barré syndrome (36 items). Initially, reporting about Guillain-Barré syndrome was more detailed; our preliminary review found cases of Guillain-Barré syndrome in eight countries in the Americas and Pacific regions, whereas microcephaly had only been reported from Brazil [19] and the first comparative study was a case-control study of Guillain-Barré syndrome [119]. Research efforts might have concentrated on congenital brain Interpretation of the review findings abnormalities since then because observations of clusters of infants with congenital abnormalities were so unusual, especially in Brazil where rubella has been eradicated. In contrast, Guillain-Barré syndrome is an established post-infectious neurological disorder and some commentators have already dubbed Zika virus “another viral cause”[15]. Our systematic approach to the assessment of causality was needed, however, because many infections have been temporally associated with Guillain-Barré symptoms [5]. Whilst the case-control study from French Polynesia is the only one published so far [119], clusters of Guillain-Barré syndrome during outbreaks of Zika virus infection have been reported from several other countries and case-control studies are ongoing in Brazil, Colombia, Mexico and Argentina.

Comparative studies based on data from the outbreak in French Polynesia suggest that the risk of both microcephaly or of Guillain-Barré syndrome is at least 30 times higher in people who had Zika virus infection compared to those who did not [54, 100, 119], although confidence intervals around these estimates are very wide. The true effect size might be weaker because the earliest studies investigating causality often overestimate the true effect, the so-called “random high” [151]. Even if the methods of other forthcoming studies in Brazil [49] and elsewhere reduce confounding and biases in selection of study populations and measurement of exposure and outcome, the increase in the risk of disease amongst those with Zika virus infection is likely to remain substantially raised. Inconsistencies in the evidence base still need investigation, however. Disease clusters have not been documented in several regions or countries affected by the most recent wave of Asian strain Zika virus infections and were not seen in Africa [152]. Some countries might not have observed these rare events because they are too small or surveillance systems are limited or use different case definitions. In the case of microcephaly, the time since the Zika outbreak might not be long enough to have resulted in births of affected babies if the period of highest risk is in the first trimester [153], or terminations of potentially affected pregnancies might have resulted in underascertainment [154].

Current evidence does not show which specific environmental and host factors interact with Zika virus to increase the risk of an affected pregnancy or of Guillain-Barré syndrome or whether there are specific factors that also have an effect in certain places. A co-factor that interacts with Zika virus to increase the risk of neurological damage could also help to explain why surveillance reports show clusters of microcephaly in some geographical areas but not others. Dengue virus has been suggested as a possible co-factor (or another component cause) [150] that might increase the risk of neurological outcomes. One major limitation to interpretation of data about causality and co-factors is the lack of accurate and accessible diagnostic tools, owing to the short duration of viraemia, cross-reactivity with other flaviviruses and lack of standardisation [155]. One report hypothesised that an insecticide used to treat drinking water (pyriproxyfen) could cause microcephaly due to possible biochemical interactions with growth regulators and observed that microcephaly cases in Brazil were reported after the introduction of the insecticide [156], but did not provide any specific data about exposure in affected women and was therefore excluded from the review.

### Strengths and limitations

The strengths of our study are that we appraised evidence of causality systematically but rapidly and transparently within a structured framework. Our searches used simple search strings and we searched sources of both published and unpublished articles without language restrictions so we believe that we identified all major relevant items about Zika virus infection. The systematic review process could not eliminate publication bias but reduced the risk that only positive reports in favour of causation would be evaluated. There were limitations to the process too, mostly resulting from the urgency of the situation. Our search strategy only included terms for Zika virus so did not cover the literature about analogous conditions or co-factors systematically. We did not have time for study selection and data extraction by two independent reviewers but additional reviewers checked the extracted data independently. Our rapid assessment of quality was not quantitative. We did not find a tool that covered our review questions and all the study designs appropriately. We followed suggestions for use of the Grading of Research Assessment Development and Evaluation tool in urgent public health situations [27] but could not standardise it for the wide range of study designs in our causality framework in the time available and we did not assign a level of certainty through formal up- or downgrading of the evidence.

### Implications for policy and research

The conclusions of the expert panel facilitate the promotion of stronger public health measures and research to tackle Zika virus and its effects. The gaps in the causality framework that we identified provide researchers with research questions and WHO has published a Zika strategic response plan [157]. Better tests to diagnose both acute and past infection will allow more accurate ascertainment of the presence of Zika virus in tissues and assessment of population level immunity to improve understanding of the epidemiology of neurological disorders. Clinical and basic research are needed to define the mechanisms of causality and to distinguish between the roles of autoimmunity and direct neurotropic effects of Zika virus in the manifestations of acute flaccid paralysis. Basic research will also further the development of vaccines, treatments and better vector control methods, which will allow direct assessment of the effects on neurological disorders of preventing Zika virus infection. For the populations currently at risk, cohort studies are needed to determine both absolute and relative risks of pregnancies affected by asymptomatic and symptomatic Zika virus infection, the role of co-factors and effect modifiers, and to define the congenital Zika virus syndrome.

### From rapid systematic review to living systematic review

Our systematic review within the structured causality framework deals with multiple neurological disorders and more detailed questions about causality than other reviews. We reached the same conclusion as Rasmussen et al. [14] but found a larger number of studies, which allowed a more comprehensive and balanced summary of evidence and of evidence gaps. In addition, our review addresses the association between Zika virus and Guillain-Barré syndrome, which is also an important source of morbidity. Systematic reviews have been done to respond to other urgent public health situations, including severe acute respiratory syndrome (SARS) [158] and avian influenza A (H5N1) [159]. These reviews focused on single conditions and only on the effects of drug treatment for which review tools are readily available.

Our review will quickly become outdated because the pace of new publications is outstripping the time taken for the review process, which includes data cleaning, analysis and interpretation. The concept of a “living systematic review” has been proposed as a way to combine rigour with timeliness for intervention research [160]. Elliott et al. propose the development of methods to produce high-quality evidence summaries that incorporate new evidence as soon as it is available and make them available immediately. The concept capitalises on technological advances to make searches and data extraction more efficient and to link the updated review text directly to open access publication with post-publication peer review [161]. The declaration by journal editors to improve access to data during public health emergencies [24, 162] could be combined with the living systematic review approach to improve the timeliness of communication about and accessibility of research about causality. We are working on methods to produce a living systematic review of the Zika causality framework that will incorporate cumulative meta-analyses of both aggregate and independent patient data as these become available.

In summary, rapid and systematic reviews with frequent updating and open dissemination are now needed, both for appraisal of the evidence about Zika virus infection and for the next public health threats that will emerge. This rapid systematic review found sufficient evidence to conclude that Zika virus is a cause of congenital abnormalities and is a trigger of Guillain-Barré situation.

## Acknowledgements

Expert panel members: Nahida Chakhtoura, National Institutes of Health/National Institute of Child Health and Development, USA; Niklas Danielsson, European Centre for Disease Prevention and Control, Sweden; Paul Garner, Liverpool School of Tropical Medicine, UK; Eva Harris, Sustainable Sciences Institute, USA; Mauricio Hernandez Avila, Instituto Nacional de Salud Pública, Mexico; Margaret Honein, Centers for Disease Control and Prevention, USA; Bart Jacobs, Erasmus University, Netherlands; Thomas Jänisch, University of Heidelberg, Germany; Marcela María Mercado Reyes, Instituto Nacional de Salud, Colombia; Ashraf Nabhan, Ain Shams University, Egypt; Laura Rodrigues, London School of Hygiene and Tropical Medicine, UK; Holger Schünemann, McMaster University, Canada; James Sejvar, Centers for Disease Control and Prevention, USA; Tom Solomon, University of Liverpool, UK; Jan Vandenbroucke, Leiden University Medical Centre, Netherlands; Vanessa Van der Linden, Hospital Barao de Lucena, Brazil; Maria van Kerkhove, Institut Pasteur, France; Tatjana Avšič Županc, University of Ljubljana, Slovenia.

WHO Zika causality working group members: Nathalie Broutet, Maria Almiron, Tarun Dua, Christopher Dye, Pierre Formenty, Florence Fouque, Metin Gulmezoglu, Edna Kara, Anais Legand, Bernadette Murgue, Susan Norris, Olufemi Oladapo, William Perea Caro, Pilar Ramon Pardo, Ludovic Reveiz Herault, João Paulo Souza.

The views expressed in this article are those of the authors and do not necessarily represent the decisions, policies, or views of the WHO or PAHO.

Supported by funding from WHO.

## Supporting information captions

**As separate files**

**S1 Text.** Background to assessment of causality in epidemiology, Zika causality framework questions, supplementary methods and results. Includes S1 Table, S2 Table, S3 Table, S1 Figure, S2 Figure, S3 Figure

**S4 Table**. S4a Table, causality framework evidence for congenital brain abnormalities; S4b Table, quality assessment of evidence about congenital brain abnormalities

**S5 Table**. S5a Table, causality framework evidence for Guillain-Barré syndrome; S5b Table, quality assessment, body of evidence

## References

1. Fauci AS, Morens DM. Zika Virus in the Americas--Yet Another Arbovirus Threat. N Engl J Med. 2016;374(7): 601–4. Epub 2016/01/14. doi: 10.1056/NEJMp1600297. PubMed PMID: 26761185.

2. Pan American Health Organization, World Health Organization. Epidemiological Alert. Neurological syndrome, congenital malformations, and Zika virus infection. Implications for public health in the Americas - 1 December 2015. 2015. Available from: http://www.paho.org/hq/index.php?option=com_docman&task=doc_view&Itemid=270&gid=32405&lang=en. [Last accessed 17.08.2016].

3. Heymann DL, Hodgson A, Sall AA, Freedman DO, Staples JE, Althabe F, et al. Zika virus and microcephaly: why is this situation a PHEIC? Lancet. 2016;387(10020): 719–21. Epub 2016/02/16. doi: 10.1016/s0140-6736(16)00320-2. PubMed PMID: 26876373.

4. Woods CG, Parker A. Investigating microcephaly. Arch Dis Child. 2013;98(9): 707–13. doi: 10.1136/archdischild-2012-302882. PubMed PMID: 23814088.

5. Willison HJ, Jacobs BC, van Doorn PA. Guillain-Barre syndrome. Lancet. 2016. doi: 10.1016/S0140-6736(16)00339-1. PubMed PMID: 26948435.

6. World Health Organization. Zika situation report. Zika virus, Microcephaly and Guillain-Barre syndrome - 4 August 2016. Geneva: World Health Organization, 2016. Available from: http://www.who.int/emergencies/zika-virus/situation-report/4-august-2016/en/. [Last accessed 10.08.2016].

7. Carey JC, Martinez L, Balken E, Leen-Mitchell M, Robertson J. Determination of human teratogenicity by the astute clinician method: review of illustrative agents and a proposal of guidelines. Birth Defects Res A Clin Mol Teratol. 2009;85(1): 63–8. doi: 10.1002/bdra.20533. PubMed PMID: 19107954.

8. Vandenbroucke JP. Observational research, randomised trials, and two views of medical science. PLoSMed. 2008;5(3):e67. doi: 10.1371/journal.pmed.0050067. PubMed PMID: 18336067; PubMed Central PMCID: PMCPMC2265762.

9. Vandenbroucke JP. In defense of case reports and case series. Ann Intern Med. 2001;134(4): 330–4. PubMed PMID: 11182844.

10. Hill AB. The Environment and Disease: Association or Causation? Proc R Soc Med. 1965;58: 295–300. PubMed PMID: 14283879; PubMed Central PMCID: PMCPMC1898525.

11. Gordis L. Chapter 14. From Association to Causation: Deriving Inferences from Epidemiologic Studies. Epidemiology: Saunders Elsevier; 2009. p.227–246.

12. Doshi P. Convicting Zika. BMJ. 2016;353: 1847. Epub 2016/04/09. doi: 10.1136/bmj.i1847. PubMed PMID: 27056643.

13. Frank C, Faber M, Stark K. Causal or not: applying the Bradford Hill aspects of evidence to the association between Zika virus and microcephaly. EMBO Mol Med. 2016;8(4): 305–7. Epub 2016/03/16. doi: 10.15252/emmm.201506058. PubMed PMID: 26976611; PubMed Central PMCID: PMCPMC4818755.

14. Rasmussen SA, Jamieson DJ, Honein MA, Petersen LR. Zika Virus and Birth Defects--Reviewing the Evidence for Causality. N Engl J Med. 2016;374(20): 1981–7. Epub 2016/04/14. doi: 10.1056/NEJMsr1604338. PubMed PMID: 27074377.

15. Smith DW, Mackenzie J.Zika virus and Guillain-Barre syndrome: another viral cause to add to the list. Lancet. 2016;387(10027): 1486–8. Epub 2016/03/08. doi: 10.1016/S0140-6736(16)00564-X. PubMed PMID: 26948432.

16. World Health Organization. Zika situation report. Zika virus, Microcephaly and Guillain-Barre syndrome - 31 March 2016. 2016. Available from: http://www.who.int/emergencies/zika-virus/situation-report/31-march-2016/en/. [Last accessed 17.08.2016].

17. Khangura S, Konnyu K, Cushman R, Grimshaw J, Moher D. Evidence summaries: the evolution of a rapid review approach. Syst Rev. 2012;1:10. doi: 10.1186/2046-4053-1-10. PubMed PMID:22587960; PubMed Central PMCID: PMCPMC3351736.

18. Egger M, Davey Smith G. Principles of and procedures for systematic reviews. In: Egger M, Davey Smith G, Altman DG, editors. Systematic Reviews in Health Care: Meta-analysis in Context. 2. London: BMJ Books; 2001. p. 23–42.

19. Broutet N, Krauer F, Riesen M, Khalakdina A, Almiron M, Aldighieri S, et al. Zika Virus as a Cause of Neurologic Disorders. N Engl J Med.2016;374(16): 1506–9. Epub 2016/03/10. doi: 10.1056/NEJMp1602708. PubMed PMID: 26959308.

20. Solomon T. Flavivirus encephalitis and other neurological syndromes (Japanese encephalitis, WNV, Tick borne encephalits, Dengue, Zika virus). Int J Infect Dis. 2016,’45:24. doi: 10.1016/j.ijid.2016.02.086. PubMed PMID: 72245053.

21. Low N, Krauer F, Riesen M. Causality framework for Zika virus and neurological disorders: systematic review protocol. PROSPERO. 2016:CRD42016036693.

22. Moher D, Shamseer L, Clarke M, Ghersi D, Liberati A, Petticrew M, et al. Preferred reporting items for systematic review and meta-analysis protocols (PRISMA-P) 2015 statement. Syst Rev. 2015;4(1): 1. doi: 10.1186/2046-4053-4-1. PubMed PMID: 25554246; PubMed Central PMCID: PMCPMC4320440.

23. Moher D, Liberati A, Tetzlaff J, Altman DG, Group P. Preferred reporting items for systematic reviews and meta-analyses: the PRISMA statement. PLoS Med. 2009;6(7):e1000097. doi: 10.1371/journal.pmed.1000097. PubMed PMID: 19621072; PubMed Central PMCID: PMCPMC2707599.

24. Dye C, Bartolomeos K, Moorthy V, Kieny MP. Data sharing in public health emergencies: a call to researchers. Bull World Health Organ. 2016;94(3): 158. doi: 10.2471/BLT.16.170860. PubMed PMID: 26966322; PubMed Central PMCID: PMCPMC4773943.

25. Zika Experimental Science Team. ZIKV-003 and ZIKV-005: Infection with French Polynesian and Asian lineage Zika virus during the first pregnancy trimester 2016 [10.05.2016]. Available from: https://zika.labkey.com/project/OConnor/ZIKV-003/begin.view?

26. National Institute for Health and Clinical Excellence. Public Health Guidance. Methods manual, Version 1: December 2005. Available from: www.nice.org.uk. London National Institute for Clinical Excellence, 2005. Available from.

27. Thayer KA, Schunemann HJ. Using GRADE to respond to health questions with different levels of urgency. Environ Int. 2016,’92-93:585–9. doi: 10.1016/j.envint.2016.03.027. PubMed PMID: 27126781.

28. Dick GW, Kitchen SF, Haddow AJ. Zika virus. I. Isolations and serological specificity. Trans R Soc Trop Med Hyg. 1952;46(5):509–20. Epub 1952/09/01. PubMed PMID: 12995440.

29. Dick GW. Zika virus. II. Pathogenicity and physical properties. Trans R Soc Trop Med Hyg. 1952;46(5):521–34. Epub 1952/09/01. PubMed PMID: 12995441.

30. Bearcroft WG. Zika virus infection experimentally induced in a human volunteer. Trans R Soc Trop Med Hyg. 1956;50(5):442–8. Epub 1956/09/01. PubMed PMID: 13380987.

31. Reagan RL, Stewart MT, Delaha EC, Brueckner AL. Response of the Syrian hamster to eleven tropical viruses by various routes of exposure. Tex Rep Biol Med. 1954;12(3): 524–7. PubMed PMID: 0007368163.

32. Weinbren MP, Williams MC. Zika virus: further isolations in the Zika area, and some studies on the strains isolated. Trans R Soc Trop Med Hyg. 1958;52(3):263–8. Epub 1958/05/01. PubMed PMID: 13556872.

33. Andral L, Bres, P., Serie, C. Yellow fever in Ethiopia. III. Serological and virological study of the forest fauna. [French]. Bull. 1968;Wld. Hlth. Org. 38(6):855–861. PubMed PMID: 0008222620.

34. Bell TM, Field EJ, Narang HK. Zika virus infection of the central nervous system of mice. Arch Gesamte Virusforsch. 1971;35(2):183–93. Epub 1971/01/01. PubMed PMID: 5002906.

35. Way JH, Bowen ET, Platt GS. Comparative studies of some African arboviruses in cell culture and in mice. J Gen Virol. 1976;30(1):123–30. Epub 1976/01/01. doi: 10.1099/0022-1317-30-1-123. PubMed PMID: 1245842.

36. Hamel R, Dejarnac O, Wichit S, Ekchariyawat P, Neyret A, Luplertlop N, et al. Biology of Zika Virus Infection in Human Skin Cells. J Virol. 2015;89(17):8880–96. Epub 2015/06/19. doi: 10.1128/JVI.00354-15. PubMed PMID: 26085147; PubMed Central PMCID: PMCPMC4524089.

37. Oliveira Melo AS, Malinger G, Ximenes R, Szejnfeld PO, Alves Sampaio S, Bispo de Filippis AM. Zika virus intrauterine infection causes fetal brain abnormality and microcephaly: tip of the iceberg? Ultrasound Obstet Gynecol. 2016;47(1):6–7. Epub 2016/01/06. doi: 10.1002/uog.15831. PubMed PMID: 26731034.

38. Ventura CV, Maia M, Bravo-Filho V, Gois AL, Belfort R, Jr. Zika virus in Brazil and macular atrophy in a child with microcephaly. Lancet. 2016;387(10015):228. Epub 2016/01/18. doi: 10.1016/S0140-6736(16)00006-4. PubMed PMID: 26775125.

39. Schuler-Faccini L, Ribeiro EM, Feitosa IM, Horovitz DD, Cavalcanti DP, Pessoa A, et al. Possible Association Between Zika Virus Infection and Microcephaly - Brazil, 2015. MMWR Morb Mortal Wkly Rep. 2016;65(3):59–62. Epub 2016/01/29. doi: 10.15585/mmwr.mm6503e2. PubMed PMID: 26820244.

40. Ventura CV, Maia M, Ventura BV, Linden VV, Araujo EB, Ramos RC, et al. Ophthalmological findings in infants with microcephaly and presumable intra-uterus Zika virus infection. Arq Bras Oftalmol. 2016;79(1):1–3. Epub 2016/02/04. doi: 10.5935/0004-2749.20160002. PubMed PMID: 26840156.

41. Mlakar J, Korva M, Tul N, Popovic M, Poljsak-Prijatelj M, Mraz J, et al. Zika Virus Associated with Microcephaly. N Engl J Med. 2016;374(10):951–8. Epub 2016/02/11. doi: 10.1056/NEJMoa1600651. PubMed PMID: 26862926.

42. de Paula Freitas B, de Oliveira Dias JR, Prazeres J, Sacramento GA, Ko AI, Maia M, et al. Ocular Findings in Infants With Microcephaly Associated With Presumed Zika Virus Congenital Infection in Salvador, Brazil. JAMA Ophthalmol. 2016. Epub 2016/02/13. doi: 10.1001/jamaophthalmol.2016.0267. PubMed PMID: 26865554.

43. Martines RB, Bhatnagar J, Keating MK, Silva-Flannery L, Muehlenbachs A, Gary J, et al. Notes from the Field: Evidence of Zika Virus Infection in Brain and Placental Tissues from Two Congenitally Infected Newborns and Two Fetal Losses--Brazil, 2015. MMWR Morb Mortal Wkly Rep. 2016;65(6):159–60. Epub 2016/02/20. doi: 10.15585/mmwr.mm6506e1. PubMed PMID: 26890059.

44. Calvet G, Aguiar RS, Melo AS, Sampaio SA, de Filippis I, Fabri A, et al. Detection and sequencing of Zika virus from amniotic fluid of fetuses with microcephaly in Brazil: a case study. Lancet Infect Dis. 2016;16(6):653–60. Epub 2016/02/22. doi: 10.1016/S1473-3099(16)00095-5. PubMed PMID: 26897108.

45. Sarno M, Sacramento GA, Khouri R, do Rosario MS, Costa F, Archanjo G, et al. Zika Virus Infection and Stillbirths: A Case of Hydrops Fetalis, Hydranencephaly and Fetal Demise. PLoS Negl Trop Dis. 2016;10(2):e0004517. Epub 2016/02/26. doi: 10.1371/journal.pntd.0004517. PubMed PMID: 26914330; PubMed Central PMCID: PMCPMC4767410.

46. Werner H, Fazecas T, Guedes B, Lopes Dos Santos J, Daltro P, Tonni G, et al. Intrauterine Zika virus infection and microcephaly: correlation of perinatal imaging and three-dimensional virtual physical models. Ultrasound Obstet Gynecol. 2016;47(5):657–60. Epub 2016/03/01. doi: 10.1002/uog.15901. PubMed PMID: 26923098.

47. Meaney-Delman D, Hills SL, Williams C, Galang RR, Iyengar P, Hennenfent AK, et al. Zika Virus Infection Among U.S. Pregnant Travelers - August 2015 - February 2016. MMWR Morb Mortal Wkly Rep. 2016;65(8):211–4. Epub 2016/03/05. doi: 10.15585/mmwr.mm6508e1. PubMed PMID: 26938703.

48. Jouannic JM, Friszer S, Leparc-Goffart I, Garel C, Eyrolle-Guignot D. Zika virus infection in French Polynesia. Lancet. 2016;387(10023):1051–2. Epub 2016/03/06. doi: 10.1016/S0140-6736(16)00625-5. PubMed PMID: 26944027.

49. Brasil P, Pereira JP, Jr., Raja Gabaglia C, Damasceno L, Wakimoto M, Ribeiro Nogueira RM, et al. Zika Virus Infection in Pregnant Women in Rio de Janeiro - Preliminary Report. N Engl J Med. 2016. Epub 2016/03/05. doi: 10.1056/NEJMoa1602412. PubMed PMID: 26943629.

50. Miranda-Filho Dde B, Martelli CM, Ximenes RA, Araujo TV, Rocha MA, Ramos RC, et al. Initial Description of the Presumed Congenital Zika Syndrome. Am J Public Health. 2016;106(4):598–600. Epub 2016/03/10. doi: 10.2105/AJPH.2016.303115. PubMed PMID: 26959258.

51. Tang H, Hammack C, Ogden SC, Wen Z, Qian X, Li Y, et al. Zika Virus Infects Human Cortical Neural Progenitors and Attenuates Their Growth. Cell Stem Cell. 2016;18(5):587–90. Epub 2016/03/10. doi: 10.1016/j.stem.2016.02.016. PubMed PMID: 26952870.

52. Villamil-Gomez WE, Mendoza-Guete A, Villalobos E, Gonzalez-Arismendy E, Uribe-Garcia AM, Castellanos JE, et al. Diagnosis, management and follow-up of pregnant women with Zika virus infection: A preliminary report of the ZIKERNCOL cohort study on Sincelejo, Colombia. Travel Med Infect Dis. 2016;14(2):155–8. Epub 2016/03/11. doi: 10.1016/j.tmaid.2016.02.004. PubMed PMID: 26960750.

53. Kleber de Oliveira W, Cortez-Escalante J, De Oliveira WT, do Carmo GM, Henriques CM, Coelho GE, et al. Increase in Reported Prevalence of Microcephaly in Infants Born to Women Living in Areas with Confirmed Zika Virus Transmission During the First Trimester of Pregnancy - Brazil, 2015. MMWR Morb Mortal Wkly Rep. 2016;65(9):242–7. Epub 2016/03/11. doi: 10.15585/mmwr.mm6509e2. PubMed PMID: 26963593.

54. Cauchemez S, Besnard M, Bompard P, Dub T, Guillemette-Artur P, Eyrolle-Guignot D, et al. Association between Zika virus and microcephaly in French Polynesia 2013-15: a retrospective study. Lancet. 2016;387(10033):2125–32. Epub 2016/03/20. doi: 10.1016/S0140-6736(16)00651-6. PubMed PMID: 26993883; PubMed Central PMCID: PMCPMC4909533.

55. Deng YQ, Zhao H, Li XF, Zhang NN, Liu ZY, Jiang T, et al. Isolation, identification and genomic characterization of the Asian lineage Zika virus imported to China. Sci China Life Sci. 2016;59(4):428- 30. Epub 2016/03/20. doi: 10.1007/s11427-016-5043-4. PubMed PMID: 26993654.

56. Faria NR, Azevedo Rdo S, Kraemer MU, Souza R, Cunha MS, Hill SC, et al. Zika virus in the Americas: Early epidemiological and genetic findings. Science. 2016;352(6283):345–9. Epub 2016/03/26. doi: 10.1126/science.aaf5036. PubMed PMID: 27013429; PubMed Central PMCID: PMCPMC4918795.

57. Rossi SL, Tesh RB, Azar SR, Muruato AE, Hanley KA, Auguste AJ, et al. Characterization of a Novel Murine Model to Study Zika Virus. Am J Trop Med Hyg. 2016;94(6):1362–9. Epub 2016/03/30. doi: 10.4269/ajtmh.16-0111. PubMed PMID: 27022155; PubMed Central PMCID: PMCPMC4889758.

58. Besnard M, Eyrolle-Guignot D, Guillemette-Artur P, Lastere S, Bost-Bezeaud F, Marcelis L, et al. Congenital cerebral malformations and dysfunction in fetuses and newborns following the 2013 to 2014 Zika virus epidemic in French Polynesia. Euro Surveill. 2016;21(13). Epub 2016/03/31. doi: 10.2807/1560-7917.ES.2016.21.13.30181. PubMed PMID: 27063794.

59. Driggers RW, Ho CY, Korhonen EM, Kuivanen S, Jaaskelainen AJ, Smura T, et al. Zika Virus Infection with Prolonged Maternal Viremia and Fetal Brain Abnormalities. N Engl J Med. 2016;374(22):2142–51. Epub 2016/03/31. doi: 10.1056/NEJMoa1601824. PubMed PMID: 27028667.

60. Dowall SD, Graham VA, Rayner E, Atkinson B, Hall G, Watson RJ, et al. A Susceptible Mouse Model for Zika Virus Infection. PLoS Negl Trop Dis. 2016;10(5):e0004658. doi: 10.1371/journal.pntd.0004658. PubMed PMID: 27149521; PubMed Central PMCID: PMCPMC4858159.

61. Homan J, Malone RW, Darnell SJ, Bremel RD. Antibody mediated epitope mimicry in the pathogenesis of Zika virus related disease. bioRxiv. 2016. doi: 10.1101/044834.

62. Pan American Health Organization. Epidemiological Alert. Zika virus infection 24 March 2016. 2016. Available from: http://www.paho.org/hq/index.php?option=com_docman&task=doc_view&Itemid=270&gid=33937&lang=en. [Last accessed 17.08.2016].

63. Pan American Health Organization. Epidemiological Alert. Zika virus infection 08 April 2016. 2016. Available from: http://www.paho.org/hq/index.php?option=com_docman&task=doc_view&Itemid=270&gid=34144&lang=en. [Last accessed 17.08.2016].

64. Pan American Health Organization. Epidemiological Update. Zika virus infection - 28 April 2016. 2016. Available from: http://www.paho.org/hq/index.php?option=com_docman&task=doc_view&Itemid=270&gid=34327&lang=en. [Last accessed 17.08.2016].

65. Pylro V, Oliveira F, Morais D, Orellana S, Pais F, Medeiros J, et al. Exploring miRNAs as the key to understand symptoms induced by ZIKA virus infection through a collaborative database. bioRxiv. 2016. doi: 10.1101/042382.

66. Dudley DM, Aliota MT, Mohr EL, Weiler AM, Lehrer-Brey G, Weisgrau KL, et al. Natural history of Asian lineage Zika virus infection in macaques. bioRxiv. 2016. doi: 10.1101/046334.

67. Reefhuis J, Gilboa SM, Johansson MA, Valencia D, Simeone RM, Hills SL, et al. Projecting Month of Birth for At-Risk Infants after Zika Virus Disease Outbreaks. Emerg Infect Dis. 2016;22(5):828–32. doi: 10.3201/eid2205.160290. PubMed PMID: 27088494; PubMed Central PMCID: PMCPMC4861542.

68. European Centre for Disease Prevention and Control. Rapid risk assessment. Zika virus disease epidemic: potential association with microcephaly and Guillain-Barre syndrome. First update 21 January 2016. 2016. Available from: http://ecdc.europa.eu/en/publications/Publications/rapid-risk-assessment-zika-virus-first-update-jan-2016.pdf. [Last accessed 17.08.2016].

69. Zmurko J, Marques RE, Schols D, Verbeken E, Kaptein SJF, Neyts J. The viral polymerase inhibitor 7-deaza-2’-C-methyladenosine is a potent inhibitor of in 1 vitro Zika virus replication and delays disease progression in a robust mouse infection model. PLoS Negl Trop Dis. 2016;10(5):no pagination. doi: 10.1101/041905.

70. Garcez PP, Loiola EC, Madeiro da Costa R, Higa LM, Trindade P, Delvecchio R, et al. Zika virus impairs growth in human neurospheres and brain organoids. Science. 2016;352(6287):816–8. Epub 2016/04/12. doi: 10.1126/science.aaf6116. PubMed PMID: 27064148.

71. Nowakowski TJ, Pollen AA, Di Lullo E, Sandoval-Espinosa C, Bershteyn M, Kriegstein AR. Expression Analysis Highlights AXL as a Candidate Zika Virus Entry Receptor in Neural Stem Cells. Cell Stem Cell. 2016;18(5):591–6. Epub 2016/04/04. doi: 10.1016/j.stem.2016.03.012. PubMed PMID: 27038591; PubMed Central PMCID: PMCPMC4860115.

72. Hazin AN, Poretti A, Turchi Martelli CM, Huisman TA, Microcephaly Epidemic Research G, Di Cavalcanti Souza Cruz D, et al. Computed Tomographic Findings in Microcephaly Associated with Zika Virus. N Engl J Med. 2016;374(22):2193–5. Epub 2016/04/07. doi: 10.1056/NEJMc1603617. PubMed PMID: 27050112.

73. World Health Organization. Zika situation report. Zika virus, Microcephaly and Guillain-Barre syndrome - 07 April 2016. 2016. Available from: http://www.who.int/emergencies/zika-virus/situation-report/7-april-2016/en/virus/situation-report/7-april-2016/en/. [Last accessed 17.08.2016].

74. Pan American Health Organization. Epidemiological Alert. Zika virus infection 31 March 2016. 2016. Available from: http://www.paho.org/hq/index.php?option=com_docman&task=doc_view&Itemid=270&gid=34041&lang=en. [Last accessed 17.08.2016].

75. Microcephaly Epidemic Research Group. Microcephaly in Infants, Pernambuco State, Brazil, 2015. Emerg Infect Dis. 2016;22(6).

76. Campanati L, Higa LM, Delvecchio R, Pezzuto P, Valadao AL, Monteiro FL, et al. The Impact of African and Brazilian ZIKV isolates on neuroprogenitors. bioRxiv. 2016. doi: 10.1101/046599.

77. Lazear HM, Govero J, Smith AM, Platt DJ, Fernandez E, Miner JJ, et al. A Mouse Model of Zika Virus Pathogenesis. Cell Host Microbe. 2016;19(5):720–30. Epub 2016/04/14. doi: 10.1016/j.chom.2016.03.010. PubMed PMID: 27066744; PubMed Central PMCID: PMCPMC4866885.

78. Bayer A, Lennemann NJ, Ouyang Y, Bramley JC, Morosky S, Marques ET, Jr., et al. Type III Interferons Produced by Human Placental Trophoblasts Confer Protection against Zika Virus Infection. Cell Host Microbe. 2016;19(5):705–12. Epub 2016/04/14. doi: 10.1016/j.chom.2016.03.008. PubMed PMID: 27066743; PubMed Central PMCID: PMCPMC4866896.

79. de Fatima Vasco Aragao M, van der Linden V, Brainer-Lima AM, Coeli RR, Rocha MA, Sobral da Silva P, et al. Clinical features and neuroimaging (CT and MRI) findings in presumed Zika virus related congenital infection and microcephaly: retrospective case series study. BMJ. 2016;353:i1901. Epub 2016/04/15. doi: 10.1136/bmj.i1901. PubMed PMID: 27075009; PubMed Central PMCID: PMCPMC4830901.

80. Cavalheiro S, Lopez A, Serra S, Da Cunha A, da Costa MD, Moron A, et al. Microcephaly and Zika virus: neonatal neuroradiological aspects. Childs Nerv Syst. 2016;32(6):1057–60. Epub 2016/04/16. doi: 10.1007/s00381-016-3074-6. PubMed PMID: 27080092; PubMed Central PMCID: PMCPMC4882355.

81. Guillemette-Artur P, Besnard M, Eyrolle-Guignot D, Jouannic JM, Garel C.Prenatal brain MRI of fetuses with Zika virus infection. Pediatr Radiol. 2016;46(7):1032–9. Epub 2016/04/20. doi: 10.1007/s00247-016-3619-6. PubMed PMID: 27090801.

82. Cordeiro MT, Pena LJ, Brito CA, Gil LH, Marques ET. Positive IgM for Zika virus in the cerebrospinal fluid of 30 neonates with microcephaly in Brazil. Lancet. 2016;387(10030):1811–2. Epub 2016/04/23. doi: 10.1016/S0140-6736(16)30253-7. PubMed PMID: 27103126.

83. Qian X, Nguyen HN, Song MM, Hadiono C, Ogden SC, Hammack C, et al. Brain-Region-Specific Organoids Using Mini-bioreactors for Modeling ZIKV Exposure. Cell. 2016;165(5):1238–54. Epub 2016/04/28. doi: 10.1016/j.cell.2016.04.032. PubMed PMID: 27118425; PubMed Central PMCID: PMCPMC4900885.

84. Paploski IA, Prates AP, Cardoso CW, Kikuti M, Silva MM, Waller LA, et al. Time Lags between Exanthematous Illness Attributed to Zika Virus, Guillain-Barre Syndrome, and Microcephaly, Salvador, Brazil. Emerg Infect Dis. 2016;22(8):1438–44. Epub 2016/05/05. doi: 10.3201/eid2208.160496. PubMed PMID: 27144515.

85. Noronha L, Zanluca C, Azevedo ML, Luz KG, Santos CN. Zika virus damages the human placental barrier and presents marked fetal neurotropism. Mem Inst Oswaldo Cruz. 2016;111(5):287- 93. Epub 2016/05/05. doi: 10.1590/0074-02760160085. PubMed PMID: 27143490; PubMed Central PMCID: PMCPMC4878297.

86. Moron AF, Cavalheiro S, Milani H, Sarmento S, Tanuri C, de Souza FF, et al. Microcephaly associated with maternal Zika virus infection. BJOG. 2016;123(8):1265–9. Epub 2016/05/07. doi: 10.1111/1471-0528.14072. PubMed PMID: 27150580.

87. Dang J, Tiwari SK, Lichinchi G, Qin Y, Patil VS, Eroshkin AM, et al. Zika Virus Depletes Neural Progenitors in Human Cerebral Organoids through Activation of the Innate Immune Receptor TLR3. Cell Stem Cell. 2016;19(2):258–65. Epub 2016/05/11. doi: 10.1016/j.stem.2016.04.014. PubMed PMID: 27162029.

88. Wu KY, Zuo GL, Li XF, Ye Q, Deng YQ, Huang XY, et al. Vertical transmission of Zika virus targeting the radial glial cells affects cortex development of offspring mice. Cell Res. 2016;26(6):645- 54. Epub 2016/05/14. doi: 10.1038/cr.2016.58. PubMed PMID: 27174054; PubMed Central PMCID: PMCPMC4897185.

89. Li C, Xu D, Ye Q, Hong S, Jiang Y, Liu X, et al. Zika Virus Disrupts Neural Progenitor Development and Leads to Microcephaly in Mice. Cell Stem Cell. 2016;19(1):120–6. Epub 2016/05/18. doi: 10.1016/j.stem.2016.04.017. PubMed PMID: 27179424.

90. Miner JJ, Cao B, Govero J, Smith AM, Fernandez E, Cabrera OH, et al. Zika Virus Infection during Pregnancy in Mice Causes Placental Damage and Fetal Demise. Cell. 2016;165(5):1081–91. Epub 2016/05/18. doi: 10.1016/j.cell.2016.05.008. PubMed PMID: 27180225; PubMed Central PMCID: PMCPMC4874881.

91. Culjat M, Darling SE, Nerurkar VR, Ching N, Kumar M, Min SK, et al. Clinical and Imaging Findings in an Infant With Zika Embryopathy. Clin Infect Dis. 2016. Epub 2016/05/20. doi: 10.1093/cid/ciw324. PubMed PMID: 27193747.

92. Johansson MA, Mier-y-Teran-Romero L, Reefhuis J, Gilboa SM, Hills SL. Zika and the Risk of Microcephaly. N Engl J Med. 2016;375(1):1–4. Epub 2016/05/26. doi: 10.1056/NEJMp1605367. PubMed PMID: 27222919; PubMed Central PMCID: PMCPMC4945401.

93. Aliota MT, Caine EA, Walker EC, Larkin KE, Camacho E, Osorio JE. Characterization of Lethal Zika Virus Infection in AG129 Mice. PLoS Negl Trop Dis. 2016;10(4):e0004682. Epub 2016/05/24. doi: 10.1371/journal.pntd.0004682. PubMed PMID: 27093158; PubMed Central PMCID: PMCPMC4836712.

94. Ventura CV, Maia M, Travassos SB, Martins TT, Patriota F, Nunes ME, et al. Risk Factors Associated With the Ophthalmoscopic Findings Identified in Infants With Presumed Zika Virus Congenital Infection. JAMA Ophthalmol. 2016. Epub 2016/05/27. doi: 10.1001/jamaophthalmol.2016.1784. PubMed PMID: 27228275.

95. Miranda HA, 2nd, Costa MC, Frazao MA, Simao N, Franchischini S, Moshfeghi DM. Expanded Spectrum of Congenital Ocular Findings in Microcephaly with Presumed Zika Infection. Ophthalmology. 2016;123(8):1788–94. Epub 2016/05/30. doi: 10.1016/j.ophtha.2016.05.001. PubMed PMID: 27236271.

96. Hanaoka MM, Kanas AF, Braconi CT, Mendes EA, Santos RA, Ferreira LCdS, et al. A Zika virus-associated microcephaly case with background exposure to STORCH agents. bioRxiv. 2016:052340. doi: 10.1101/052340.

97. Garcez PP, Nascimento JM, Mota de Vasconcelos J, Madeiro da Costa R, Delvecchio R, Trindade P, et al. Combined proteome and transcriptome analyses reveal that Zika virus circulating in Brazil alters cell cycle and neurogenic programmes in human neurospheres. PeerJ Preprints. 2016. doi:https://doi.org/10.7287/peeri.preprints.2033v1.

98. Ganguly B, Ganguly E. Disruption of human astn2 function by ZIKV ns4b gene as a molecular basis for Zika viral microcephaly. bioRxiv. 2016:054486. doi: 10.1101/054486.

99. Frumence E, Roche M, Krejbich-Trotot P, El-Kalamouni C, Nativel B, Rondeau P, et al. The South Pacific epidemic strain of Zika virus replicates efficiently in human epithelial A549 cells leading to IFN-beta production and apoptosis induction. Virology. 2016;493:217–26. doi: 10.1016/j.virol.2016.03.006. PubMed PMID: 27060565.

100. Araujo TVB, Rodrigues LC, Ximenes RA, Miranda-Filho DB, Montarroyos UR, Melo APL, et al. Microcephaly and Zika infection: Preliminary Report of a Case-Control Study. Lancet. 2016:In press.

101. Gregg NM. Congenital Cataract Following German Measles in the Mother, 1941. Epidemiol Infect. 1941;107(1):35–46.

102. Cheeran MC, Lokensgard JR, Schleiss MR. Neuropathogenesis of congenital cytomegalovirus infection: disease mechanisms and prospects for intervention. Clin Microbiol Rev. 2009;22(1):99–126, Table of Contents. doi: 10.1128/CMR.00023-08. PubMed PMID: 19136436; PubMed Central PMCID: PMCPMC2620634.

103. O’Leary DR, Kuhn S, Kniss KL, Hinckley AF, Rasmussen SA, Pape WJ, et al. Birth outcomes following West Nile Virus infection of pregnant women in the United States: 2003-2004. Pediatrics. 2006;117(3):e537–45. doi: 10.1542/peds.2005-2024. PubMed PMID: 16510632.

104. Pouliot SH, Xiong X, Harville E, Paz-Soldan V, Tomashek KM, Breart G, et al. Maternal dengue and pregnancy outcomes: a systematic review. Obstet Gynecol Surv. 2010;65(2):107–18. doi: 10.1097/OGX.0b013e3181cb8fbc. PubMed PMID: 20100360.

105. Franca GV, Schuler-Faccini L, Oliveira WK, Henriques CM, Carmo EH, Pedi VD, et al. Congenital Zika virus syndrome in Brazil: a case series of the first 1501 livebirths with complete investigation. Lancet. 2016. Epub 2016/07/04. doi: 10.1016/s0140-6736(16)30902-3. PubMed PMID: 27372398.

106. Musso DG, D.J.Zika Virus. Clin Microbiol Rev. 2016;29(3):487–524. Epub 2016/04/01. doi: 10.1128/cmr.00072-15. PubMed PMID: 27029595.

107. Duffy MR, Chen TH, Hancock WT, Powers AM, Kool JL, Lanciotti RS, et al. Zika virus outbreak on Yap Island, Federated States of Micronesia. N Engl J Med. 2009;360(24):2536–43. Epub 2009/06/12. doi: 10.1056/NEJMoa0805715. PubMed PMID: 19516034.

108. World Health Organization WHO. Zika situation report. Neurological syndrome and congenital anomalies - 5 February 2016. 2016. Available from: http://apps.who.int/iris/bitstream/10665/204348/1/zikasitrep_5Feb2016_eng.pdf?ua=1.

109. Oehler E, Watrin L, Larre P, Leparc-Goffart I, Lastere S, Valour F, et al. Zika virus infection complicated by Guillain-Barre syndrome--case report, French Polynesia, December 2013.Euro Surveill. 2014;19(9). Epub 2014/03/15. PubMed PMID: 24626205.

110. Institut de veille sanitaire. Virus Zika Polynésie 2013-2014, Ile de Yap, Micronésie 2007 - Janvier 2014. 2014. Available from: http://www.invs.sante.fr/Publications-et-outils/Points-epidemiologiques/Tous-les-numeros/International/Virus-Zika-en-Polynesie-2013-2014-et-ile-de-Yap-Micronesie-2007-Janvier-2014. [Last accessed 17.8.16].

111. Ioos S, Mallet HP, Leparc Goffart I, Gauthier V, Cardoso T, Herida M. Current Zika virus epidemiology and recent epidemics. Med Mal Infect. 2014;44(7):302–7. Epub 2014/07/09. doi: 10.1016/j.medmal.2014.04.008. PubMed PMID: 25001879.

112. Pan American Health Organization. Epidemiological Update. Neurological syndrome, congenital anomalies, and Zika virus infection - 17 January 2016. 2016. Available from: http://www.paho.org/hq/index.php?option=com_docman&task=doc_view&Itemid=270&gid=32879&lang=en. [Last accessed 17.08.2016].

113. Pan American Health Organization. Epidemiological Alert. Zika virus infection -3 March 2016. 2016. Available from: http://www.paho.org/hq/index.php?option=com_docman&task=doc_view&Itemid=270&gid=33486&lang=en. [Last accessed 17.08.2016].

114. Pan American Health Organization. Epidemiological Alert. Zika virus infection -10 March 2016. 2016. Available from: http://www.paho.org/hq/index.php?option=com_docman&task=doc_view&Itemid=270&gid=33659&lang=en. [Last accessed 17.08.2016].

115. Pan American Health Organization. Epidemiological Alert. Zika virus infection -17 March 2016. 2016. Available from: http://www.paho.org/hq/index.php?option=com_docman&task=doc_view&Itemid=270&gid=33768&lang=en. [Last accessed 16.08.2016].

116. Pan American Health Organization. Epidemiological Update. Zika virus infection - 21 April 2016. 2016. Available from: http://www.paho.org/hq/index.php?option=com_docman&task=doc_view&Itemid=270&gid=34243&lang=en. [Last accessed 17.08.2016].

117. Thomas DL, Sharp TM, Torres J, Armstrong PA, Munoz-Jordan J, Ryff KR, et al. Local Transmission of Zika Virus--Puerto Rico, November 23, 2015- January 28, 2016. MMWR Morb Mortal Wkly Rep. 2016; 65(6):154–8. Epub 2016/02/20. doi: 10.15585/mmwr.mm6506e2. PubMed PMID: 26890470.

118. Reyna-Villasmil E, Lopez-Sanchez G, Santos-Bolivar J. [Guillain-Barre syndrome due to Zika virus during pregnancy]. Med Clin (Barc). 2016;146(7):331–2. Epub 2016/03/08. doi: 10.1016/j.medcli.2016.02.002. PubMed PMID: 26947168.

119. Cao-Lormeau VM, Blake A, Mons S, Lastere S, Roche C, Vanhomwegen J, et al. Guillain-Barre Syndrome outbreak associated with Zika virus infection in French Polynesia: a case-control study. Lancet. 2016;387(10027):1531–9. Epub 2016/03/08. doi: 10.1016/S0140-6736(16)00562-6. PubMed PMID: 26948433.

120. Roze B, Najioullah F, Ferge JL, Apetse K, Brouste Y, Cesaire R, et al. Zika virus detection in urine from patients with Guillain-Barre syndrome on Martinique, January 2016. Euro Surveill. 2016;21(9). Epub 2016/03/12. doi: 10.2807/1560-7917.ES.2016.21.9.30154. PubMed PMID: 26967758.

121. Lucchese G, Kanduc D. Zika virus and autoimmunity: From microcephaly to Guillain-Barre syndrome, and beyond. Autoimmun Rev. 2016;15(8):801–8. Epub 2016/03/29. doi: 10.1016/j.autrev.2016.03.020. PubMed PMID: 27019049.

122. Craig AT, Butler MT, Pastore R, Paterson BJ, Durrheim DN. Update on Zika virus transmission in the Pacific islands, 2007 to February 2016 and failure of acute flaccid paralysis surveillance to signal Zika emergence in this setting. Bull World Health Organ. 2016. doi: 10.2471/blt.16.171892.

123. Watrin L, Ghawche F, Larre P, Neau JP, Mathis S, Fournier E. Guillain-Barre Syndrome (42 Cases) Occurring During a Zika Virus Outbreak in French Polynesia. Medicine (Baltimore). 2016;95(14):e3257. Epub 2016/04/09. doi: 10.1097/MD.0000000000003257. PubMed PMID: 27057874.

124. Fontes CA, Dos Santos AA, Marchiori E. Magnetic resonance imaging findings in Guillain-Barre syndrome caused by Zika virus infection. Neuroradiology. 2016. Epub 2016/04/14. doi: 10.1007/s00234-016-1687-9. PubMed PMID: 27067205.

125. Brasil P, Sequeira PC, Freitas AD, Zogbi HE, Calvet GA, de Souza RV, et al. Guillain-Barre syndrome associated with Zika virus infection. Lancet. 2016;387(10026):1482. Epub 2016/04/27. doi: 10.1016/S0140-6736(16)30058-7. PubMed PMID: 27115821.

126. Kassavetis P, Joseph JM, Francois R, Perloff MD, Berkowitz AL. Zika virus-associated Guillain-Barre syndrome variant in Haiti. Neurology. 2016;87(3):336–7. Epub 2016/05/11. doi: 10.1212/WNL.0000000000002759. PubMed PMID: 27164708.

127. Duijster JW, Goorhuis A, van Genderen PJ, Visser LG, Koopmans MP, Reimerink JH, et al. Zika virus infection in 18 travellers returning from Surinam and the Dominican Republic, The Netherlands, November 2015-March 2016. Infection. 2016. Epub 2016/05/23. doi: 10.1007/s15010-016-0906-y. PubMed PMID: 27209175.

128. van den Berg B, van den Beukel JC, Alsma J, van der Eijk AA, Ruts L, van Doorn PA, et al. [Guillain-Barre syndrome following infection with the Zika virus]. Ned Tijdschr Geneeskd. 2016;160(0):D155. Epub 2016/05/28. PubMed PMID: 27229696.

129. Yung CFT, K.C.Guillain-Barre Syndrome and Zika Virus: Estimating Attributable Risk to Inform Intensive Care Capacity Preparedness. Clin Infect Dis. 2016. Epub 2016/05/27. doi: 10.1093/cid/ciw355. PubMed PMID: 27225243.

130. Jackson BR, Zegarra JA, Lopez-Gatell H, Sejvar J, Arzate F, Waterman S, et al. Binational outbreak of Guillain-Barre syndrome associated with Campylobacter jejuni infection, Mexico and USA, 2011. Epidemiol Infect. 2014;142(5):1089–99. doi: 10.1017/S0950268813001908. PubMed PMID: 23924442.

131. McCarthy N, Giesecke J. Incidence of Guillain-Barre syndrome following infection with Campylobacter jejuni. Am J Epidemiol. 2001;153(6):610–4. PubMed PMID: 11257070.

132. Tam CC, Rodrigues LC, Petersen I, Islam A, Hayward A, O’Brien SJ. Incidence of Guillain-Barre syndrome among patients with Campylobacter infection: a general practice research database study. J Infect Dis. 2006;194(1):95–7. doi: 10.1086/504294. PubMed PMID: 16741887.

133. Rees JH, Soudain SE, Gregson NA, Hughes RA. Campylobacter jejuni infection and Guillain-Barre syndrome. N Engl J Med. 1995;333(21):1374–9. doi: 10.1056/NEJM199511233332102. PubMed PMID: 7477117.

134. Mishu B, Blaser MJ. Role of infection due to Campylobacter jejuni in the initiation of Guillain-Barre syndrome. Clin Infect Dis. 1993;17(1):104–8. PubMed PMID: 8353228.

135. Carod-Artal FJ, Wichmann O, Farrar J, Gascon J. Neurological complications of dengue virus infection. Lancet Neurol. 2013;12(9):906–19. Epub 2013/08/21. doi: 10.1016/S1474-4422(13)70150-9. PubMed PMID: 23948177.

136. Sejvar JJ, Bode AV, Marfin AA, Campbell GL, Ewing D, Mazowiecki M, et al. West Nile virus-associated flaccid paralysis. Emerg Infect Dis. 2005;11(7):1021–7. doi: 10.3201/eid1107.040991. PubMed PMID: 16022775; PubMed Central PMCID: PMCPMC3371783.

137. Xiang JY, Zhang YH, Tan ZR, Huang J, Zhao YW. Guillain-Barre syndrome associated with Japanese encephalitis virus infection in China. Viral Immunol. 2014;27(8):418–20. doi: 10.1089/vim.2014.0049. PubMed PMID: 25140441.

138. Ravi V, Taly AB, Shankar SK, Shenoy PK, Desai A, Nagaraja D, et al. Association of Japanese encephalitis virus infection with Guillain-Barre syndrome in endemic areas of south India. Acta Neurol Scand. 1994;90(1):67–72. PubMed PMID: 7941960.

139. McMahon AW, Eidex RB, Marfin AA, Russell M, Sejvar JJ, Markoff L, et al. Neurologic disease associated with 17D-204 yellow fever vaccination: a report of 15 cases. Vaccine. 2007;25(10):1727- 34. doi: 10.1016/j.vaccine.2006.11.027. PubMed PMID: 17240001.

140. Solomon T, Kneen R, Dung NM, Khanh VC, Thuy TT, Ha DQ, et al. Poliomyelitis-like illness due to Japanese encephalitis virus. Lancet. 1998;351(9109):1094–7. doi: 10.1016/S0140-6736(97)07509-0. PubMed PMID: 9660579.

141. Chung CC, Lee SS, Chen YS, Tsai HC, Wann SR, Kao CH, et al. Acute flaccid paralysis as an unusual presenting symptom of Japanese encephalitis: a case report and review of the literature. Infection. 2007;35(1):30–2. doi: 10.1007/s15010-007-6038-7. PubMed PMID: 17297587.

142. Argall KG, Armati PJ, King NJ, Douglas MW. The effects of West Nile virus on major histocompatibility complex class I and II molecule expression by Lewis rat Schwann cells in vitro. J Neuroimmunol. 1991;35(1-3):273–84. PubMed PMID: 1955569.

143. Loshaj-Shala A, Regazzoni L, Daci A, Orioli M, Brezovska K, Panovska AP, et al. Guillain Barre syndrome (GBS): new insights in the molecular mimicry between C.jejuni and human peripheral nerve (HPN) proteins. J Neuroimmunol. 2015;289:168–76. doi: 10.1016/j.jneuroim.2015.11.005. PubMed PMID: 26616887.

144. Meyer Sauteur PM, Huizinga R, Tio-Gillen AP, Roodbol J, Hoogenboezem T, Jacobs E, et al. Mycoplasma pneumoniae triggering the Guillain-Barre syndrome: A case-control study. Ann Neurol. 2016. doi: 10.1002/ana.24755. PubMed PMID: 27490360.

145. Tseng YF, Wang CC, Liao SK, Chuang CK, Chen WJ. Autoimmunity-related demyelination in infection by Japanese encephalitis virus. J Biomed Sci. 2011,18:20. doi: 10.1186/1423-0127-18-20. PubMed PMID: 21356046; PubMed Central PMCID: PMCPMC3056755.

146. Dejnirattisai W, Supasa P, Wongwiwat W, Rouvinski A, Barba-Spaeth G, Duangchinda T, et al. Dengue virus sero-cross-reactivity drives antibody-dependent enhancement of infection with zika virus. Nat Immunol. 2016. Epub 2016/06/25. doi: 10.1038/ni.3515. PubMed PMID: 27339099.

147. Paul LM, Carlin ER, Jenkins MM, Tan AL, Barcellona CM, Nicholson CO, et al. Dengue Virus Antibodies Enhance Zika Virus Infection. bioRxiv. 2016:050112. doi: 10.1101/050112.

148. World Health Organization. Pregnancy management in the context of Zika virus infection. Interim guidance update. Geneva: World Health Organization, 2016. Available from: http://www.who.int/csr/resources/publications/zika/pregnancy-management/en/. [Last accessed 18.08.2016].

149. Rothman KJ, Greenland S. Causation and causal inference in epidemiology. Am J Public Health. 2005;95 Suppl 1:S144-50. doi: 10.2105/AJPH.2004.059204. PubMed PMID: 16030331.

150. Rothman KJ. Causes. Am J Epidemiol. 1976;104(6):587–92. PubMed PMID: 998606.

151. Vandenbroucke JP. Case reports in an evidence-based world. J R Soc Med. 1999;92(4):159–63. PubMed PMID: 10450190; PubMed Central PMCID: PMC1297135.

152. Solomon T, Baylis M, Brown D. Zika virus and neurological disease-approaches to the unknown. Lancet Infect Dis. 2016. Epub 2016/03/01. doi: 10.1016/s1473-3099(16)00125-0. PubMed PMID: 26923117.

153. Ribeiro LS, Marques RE, Jesus AM, Almeida RP, Teixeira MM. Zika crisis in Brazil: challenges in research and development. Curr Opin Virol. 2016;18:76–81. Epub 2016/05/18. doi: 10.1016/j.coviro.2016.04.002. PubMed PMID: 27179929.

154. Aiken AR, Scott JG, Gomperts R, Trussell J, Worrell M, Aiken CE. Requests for Abortion in Latin America Related to Concern about Zika Virus Exposure. N Engl J Med. 2016;375(4):396–8. doi: 10.1056/NEJMc1605389. PubMed PMID: 27331661.

155. Haug CJ, Kieny MP, Murgue B. The Zika Challenge. N Engl J Med. 2016;374(19):1801–3. Epub 2016/03/31. doi: 10.1056/NEJMp1603734. PubMed PMID: 27028782.

156. Evans DN, Fred; Parens, Raphael; Morales, Alfredo J.; Bar-Yam, Yaneer. A Possible Link Between Pyriproxyfen and Microcephaly. arXiv. 2016.

157. Triunfol M. Microcephaly in Brazil: confidence builds in Zika connection. Lancet Infect Dis. 2016;16(5):527–8. doi: 10.1056/NEJMoa1602412. PubMed PMID: WOS:000374272900019.

158. Stockman LJ, Bellamy R, Garner P. SARS: systematic review of treatment effects. PLoS Med. 2006;3(9):e343. doi: 10.1371/journal.pmed.0030343. PubMed PMID: 16968120; PubMed Central PMCID: PMCPMC1564166.

159. Schunemann HJ, Hill SR, Kakad M, Bellamy R, Uyeki TM, Hayden FG, et al. WHO Rapid Advice Guidelines for pharmacological management of sporadic human infection with avian influenza A (H5N1) virus. Lancet Infect Dis. 2007;7(1):21–31. doi: 10.1016/S1473-3099(06)70684-3. PubMed PMID: 17182341.

160. Elliott JH, Turner T, Clavisi O, Thomas J, Higgins JPT, Mavergames C, et al. Living Systematic Reviews: An Emerging Opportunity to Narrow the Evidence-Practice Gap. PLoS Med. 2014;11(2):1–6. doi: 10.1371/journal.pmed.1001603. PubMed PMID: Elliott2014.

161. Tracz V, Lawrence R. Towards an open science publishing platform. F1000Res. 2016;5:130. doi: 10.12688/f1000research.7968.1. PubMed PMID: 26962436; PubMed Central PMCID: PMCPMC4768651.

162. Modjarrad K, Moorthy VS, Millett P, Gsell PS, Roth C, Kieny MP. Developing Global Norms for Sharing Data and Results during Public Health Emergencies. PLoS Med. 2016;13(1):e1001935. doi: 10.1371/journal.pmed.1001935. PubMed PMID: 26731342; PubMed Central PMCID: PMCPMC4701443.

